# Skeletal progenitor LRP1 deficiency causes severe and persistent skeletal defects with WNT/planar cell polarity dysregulation

**DOI:** 10.1101/2023.07.11.548556

**Authors:** Mohammad Alhashmi, Abdulrahman ME Gremida, Santosh K Maharana, Marco Antonaci, Amy Kerr, Noor A Al-Maslamani, Ke Liu, Maria M Meschis, Hazel Sutherland, Peter Wilson, Peter Clegg, Grant N Wheeler, Robert J van ’t Hof, George Bou-Gharios, Kazuhiro Yamamoto

**Affiliations:** Institute of Life Course and Medical Sciences, University of Liverpool, 6 West Derby Street, Liverpool L7 8TX, United Kingdom; School of Biological Sciences, University of East Anglia, Norwich Research Park, Norwich, Norfolk NR4 7TJ, United Kingdom

**Author notes:** **Corresponding author:** Kazuhiro Yamamoto, William Henry Duncan Building, University of Liverpool, 6 West Derby Street, Liverpool, L7 8TX, UK. These authors equally contributed to the work (Mohammad Alhashmi, Abdulrahman ME Gremida).

**Keywords:** Bone, synovial joint, endocytosis, perichondrium, hip dysplasia, osteoarthritis, Wnt, Planar cell polarity, Xenopus

## Abstract

Low-density lipoprotein receptor-related protein 1 (LRP1) is a multifunctional endocytic receptor whose dysfunction is linked to developmental dysplasia of the hip, osteoporosis and osteoarthritis. Our work addresses the critical question of how these skeletal pathologies emerge. Here, we show the abundant expression of LRP1 in skeletal progenitor cells at mouse embryonic stage E13.5 and onwards, especially in the perichondrium, the stem cell layer surrounding developing limbs essential for bone formation. *Lrp1* deficiency in these stem cells causes joint fusion, malformation of cartilage/bone template and markedly delayed or lack of primary ossification along with aberrant accumulation of some of the LRP1 ligands at as early as E16.5. These early abnormalities result in multiple and persistent skeletal defects including a severe deficit in hip joint and patella, and markedly deformed and low-density long bones leading to dwarfism and impaired mobility. Mechanistically, we show that LRP1 regulates core non-canonical WNT/planar cell polarity (PCP) components that may explain the malformation of long bones. LRP1 directly binds to Wnt5a, facilitates its cell-association and endocytic recycling. Using *Xenopus* as a model system we show that loss or gain of LRP1 function leads to shortened tadpoles similar to Wnt5a and Wnt11 overexpression, indicating a role for LRP1 in WNT/PCP signalling. Finally, we show the colocalisation LRP1 and Wnt5a in the developing mouse limbs and that *Lrp1* deficiency diminishes graded distribution of Wnt5a and Vangl2. We propose that skeletal progenitor LRP1 plays a critical role in formation and maturity of multiple bones and joints by regulating morphogen signalling, providing novel insights into the fundamental processes of morphogenesis and the emergence of skeletal pathologies.

## INTRODUCTION

The low-density lipoprotein (LDL) receptor-related protein 1 (LRP1) is widely expressed type 1 transmembrane protein in adult tissues (1, 2) that regulates cellular events by modulating the levels of structurally and functionally diverse extracellular molecules via clathrin-dependent endocytosis (3, 4). Our previous studies demonstrated that LRP1 plays an important role in the turnover of extracellular matrix (ECM) components in articular cartilage by mediating endocytic clearance of cartilage-degrading proteinases and their inhibitors (5–10). This endocytic process is impaired in cartilage under inflammatory conditions or in osteoarthritis (OA), the most prevalent age-related degenerative joint disease (11). LRP1 also participates in signalling pathways through interaction with membrane receptors and cytoplasmic adaptor proteins. The cytoplasmic NPxY motifs within the LRP1 intracellular domain also provide binding sites for a set of signalling proteins (12–14).

Global deletion of the *Lrp1* gene in mice results in early embryonic lethality at E13.5 (15, 16). The homozygous and heterozygous *Lrp1* null^tm1.1(KOMP)Wtsi^ mice exhibit multiple phenotypes, as described in the Mouse Genome Informatics database (MGI: 5495233). Tissue-specific *Lrp1* deletion in mice has revealed various biological roles of LRP1 including lipoprotein metabolism (17, 18), insulin signalling (19), inflammation (20, 21), heart development (22), vascular wall integrity and remodelling (23–26) and bone development and remodelling (27–30). Together, these studies highlight a critical and non-redundant role of LRP1 in development as well as adult tissue homeostasis.

*LRP1* single nucleotide polymorphisms are associated with a decrease in bone mineral density and content (31). A recent study by Yan *et al* (32) identified mutations in *LRP1* including R1783W with developmental dysplasia of the hip (DDH) patients. In mice, mutant *Lrp1^R1783W^*homozygote and heterozygote mice exhibited delayed Y-shaped triradiate cartilage and smaller acetabulum in 8-week and 16-week-old mice, respectively (32). Heterozygous global *Lrp1* knockout (KO) mice also developed a hip dysplasia phenotype. In contrast, a bone and cartilage conditional KO mice (*Lrp1*^flox/flox^*/Col2a1*^Cre^) showed shortened bones and cartilage growth plate (27). In vitro, siRNA-mediated Lrp1 gene-silencing reduced chondrogenesis of human mesenchymal stem cells (33). These studies suggest that function of LRP1 during synovial joint and bone formation is cell type- and time-dependent. However, LRP1 expression and distribution in the developmental stages remains incompletely understood.

In this study, we addressed the critical questions of when and where LRP1 is expressed during skeletal development, how deficiency of LRP1 leads to skeletal pathologies and how long it persists, and which molecular mechanisms underpin the defects. We showed that LRP1 is abundantly expressed in skeletal progenitor cells at embryonic stage E13.5 and onwards, in particular in the perichondrium (34). To investigate the role of skeletal progenitor LRP1, we generated a conditional KO of *Lrp1* using paired-related homebox gene-1 (*Prrx1*), which is expressed in the early limb bud skeletal mesenchyme and a subset of craniofacial mesenchyme (35, 36). *Lrp1*^flox/flox^*/Prrx1*^Cre^ mice exhibited severe and persistent malformation of multiple bones and synovial joints, which were not evident in either the *Lrp1^R1783W^* or *Lrp1*^flox/flox^*/Col2a1*^Cre^ mice. Our exploration of molecular mechanisms showed unique regulation of non-canonical WNT/planar cell polarity (PCP) components and signalling by LRP1. We propose that LRP1 plays a critical role in skeletal development by regulating WNT morphogen signalling, which governs a myriad of biological processes underlying the development and maintenance of adult tissue homeostasis.

## RESULTS

### LRP1 is abundantly expressed in skeletal progenitor cells, in particular in the perichondrium

We investigated the distribution of LRP1 during skeletal development, which had thus far remained unknown. Histological investigation of LRP1 protein in developing limbs showed that LRP1 is abundantly expressed in E13.5- to newborn (P0) elbow joints, with the strongest immunosignals of LRP1 detected in the perichondrium layers, a dense stem cell layer surrounding developing limbs essential for bone formation (**Fig 1A**). To investigate the role of skeletal progenitor LRP1 in skeletal development *in vivo*, we have established a double transgenic mouse line *Lrp1*^flox/flox^*/Prrx1*^Cre^ (**Fig 1B**). In this strain, a 2.4 kb *Prrx1* enhancer directs the transgene expression in undifferentiated mesenchyme in the developing limb buds around embryonic day 9.5 (E9.5) (35–38). Transgene expression is extinguished in the condensing mesenchyme and chondrocytes, and the expression is confined to the perichondrium/periosteum of the limbs at E15.5. Immunohistochemistry of LRP1 confirmed specific deletion of LRP1 in *Prrx1* expressing cells in the E16.5 knee but not in their ribs (**Fig 1C**).

**Fig 1.**
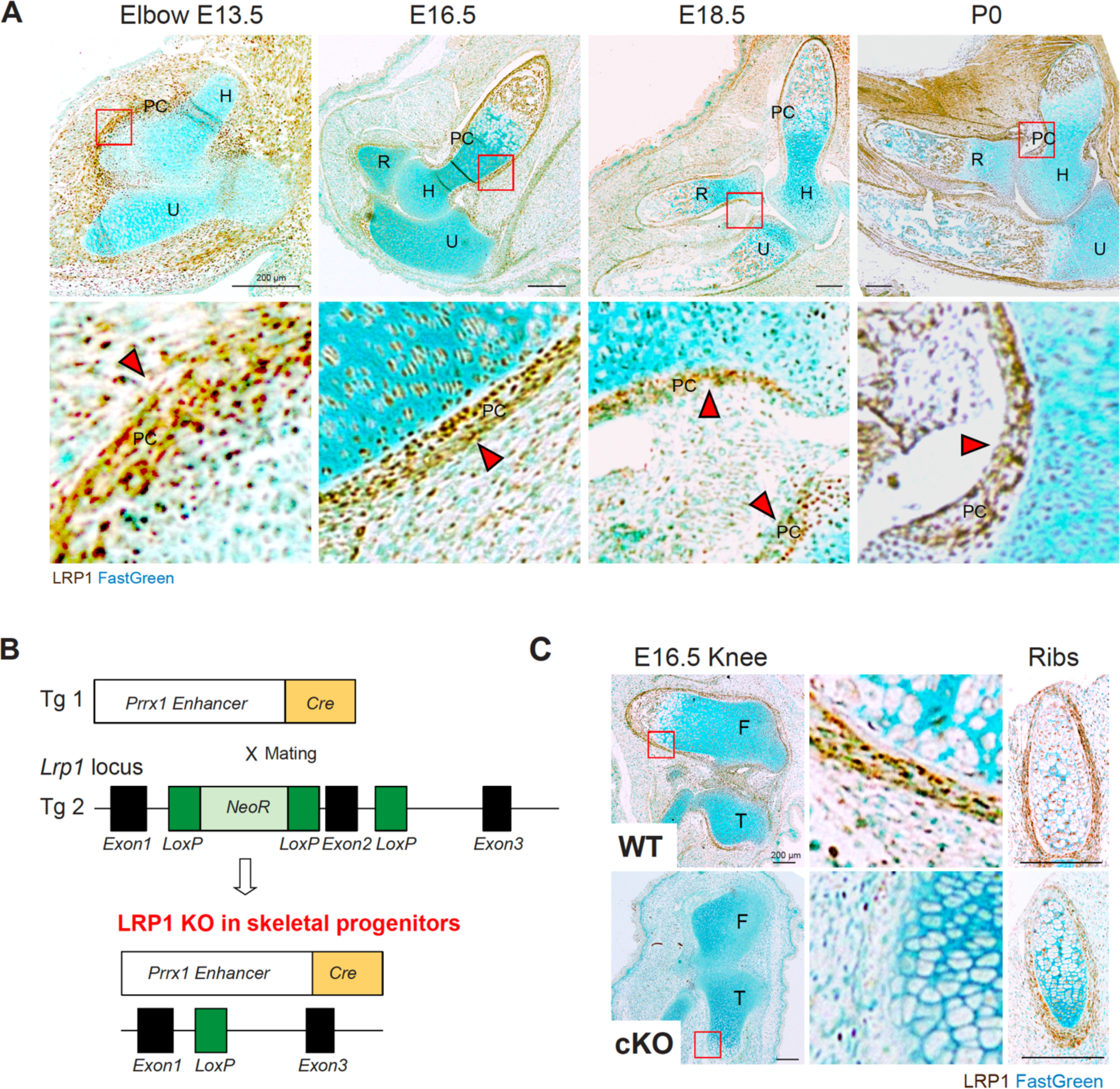
LRP1 is abundantly expressed in skeletal progenitor cells, in particular in the perichondrium. *A*, Representative images of immunohistochemical staining of LRP1 and fast green counterstaining in WT E13.5-P0 elbow joint sections of *Lrp1*^flox/flox^ mice. Scale bar, 200 µm. H, humerus; R, radius; U, ulna; PC, perichondrium. Regions delineated by the red squares in the panels have been magnified in the lower panels. Arrow heads indicate abundant LRP1 expression in perichondrium layers. *B*, Schematic diagram showing the constructs used to generate skeletal progenitor-selective LRP1 conditional knockout mice. Transgenic mouse lines harbouring Prrx1 limb enhancer Cre (Tg 1) and floxed *LRP1* (Tg 2) were used to establish the LRP1Prrx1Cre line. *C*, Representative images of immunohistochemical staining of LRP1 and fast green counterstaining in E16.5 knee and rib sections of WT and *Lrp1*^flox/flox^*/Prrx1*^Cre^ homozygote conditional KO (cKO) mice. Regions delineated by the red squares in the upper panels have been magnified in the lower panels. Scale bar, 200 µm. F, femur; T, tibia.

### Conditional deletion of *Lrp1* in skeletal progenitors impairs early bone and joint formation

Haematoxylin and eosin (H&E) staining of limbs at different embryonic stages revealed fusion of joints, malformation of cartilage/bone template and markedly delayed or lack of primary ossification in E16.5 *Lrp1*^flox/flox^*/Prrx1*^Cre^ homozygote shoulder, elbow and knee joints (**Fig 2A-D**) and the defects become more severe in E18.5 and P0 neonates (**Fig 2BC**). Striking joint malformations were observed in P0 *Lrp1*^flox/flox^*/Prrx1*^Cre^ hip joints (**Fig 2D**). These defects were not observed in heterozygote (*Lrp1*^flox/wt^*/Prrx1*^Cre^) littermates. Histological investigation of LRP1 ligands including tissue inhibitor of matrix metalloproteinase 3 (TIMP3) and CCN2 revealed their aberrant accumulation in P0 *Lrp1*^flox/flox^*/Prrx1*^Cre^ compared with WT limbs (**Fig 2HI**), indicating a critical role of LRP1 in their tissue availability. High resolution µCT scanning showed that femur and humerus length were shorter in P0 *Lrp1*^flox/flox^*/Prrx1*^Cre^ compared with WT mice, whereas no significant difference was observed in E18.5 embryos (**Fig 2J**). No significant differences in body length were observed in P0 neonates (**Fig 2K**).

**Fig 2.**
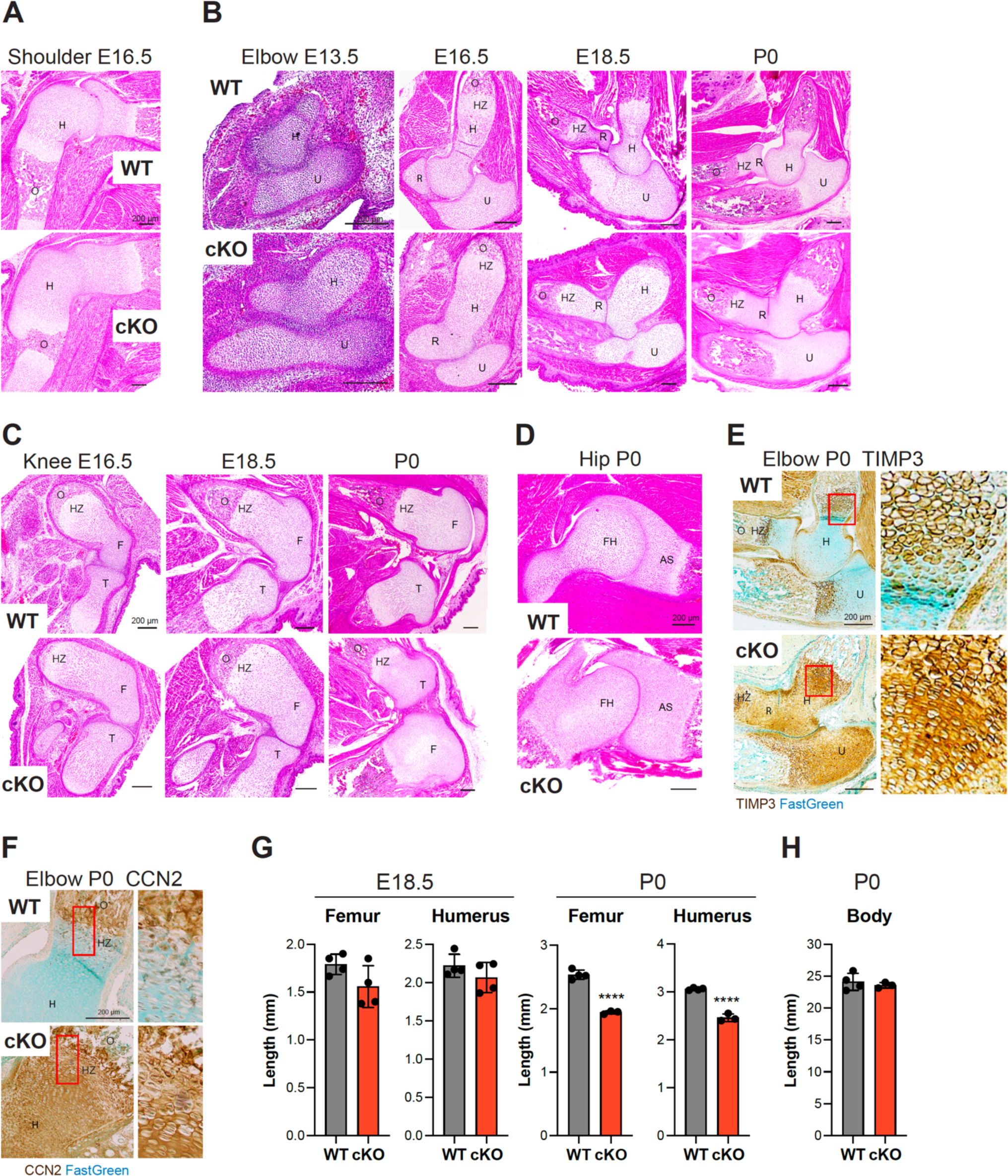
Conditional deletion of *Lrp1* in skeletal progenitors impairs early bone and joint formation. *A-D*, Representative images of H&E staining of E16.5 shoulder (*A*), E13.5-P0 elbow (*B*), E16.5-P0 knee (*C*) and P0 hip (*D*) sections of WT and *Lrp1*^flox/flox^*/Prrx1*^Cre^ homozygote conditional KO (cKO) mice. Scale bar, 200 µm. H, humerus; R, radius; U, ulna; HZ, hypertrophic zone; O, primary ossification center; F, femur; FH, femur head; T, tibia; AS, acetabulum socket. *E* and *F*, Representative images of immunohistochemical staining of TIMP3 (*E*) and CCN2 (*F*) and fast green counterstaining in E16.5 knee sections of WT and cKO mice. Regions delineated by the red squares in the panels have been magnified in the right panels. Scale bar, 200 µm. *G* and *H*, Length of femur and humerus bone (*G*) and body (*H*) of E18.5 (*G*) and P0 (*G* and *H*) WT and cKO mice. Circles represent individual mice and bars show the mean ± *SD*. *p* values were evaluated by 2-tailed Student’s t test. ****, *p* < 0.0001.

These joint and cartilage/bone template defects have not been reported for the *Lrp1^R1783W^* (32) or *Lrp1*^flox/flox^*/Col2a1*^Cre^ mice (27) most likely because they reflect events that occurred earlier than *Col2a1* transcriptional activation. To further evaluate the role of chondrocyte LRP1 in the cKO early skeletal phenotype, we generated a double transgenic mouse line *Lrp1*^flox/flox^*/Acan*^CreERT2^ (39). In contrast to *Prrx1* expression (36, 37), *Aggrecan* expression was detected in the E13.0 forelimb and in all developing cartilage by E15.5 (39). *Lrp1* was deleted at the different embryonic stages and E19.5 embryos were examined (**Fig S1**). *Lrp1* gene excision was confirmed by *Lrp1 exon 2* PCR (**Fig S1B**) but histological analysis and bone length measurement confirmed normality of skeletal development in E19.5 *Lrp1*^flox/flox^*/Acan*^CreERT2^ mice (**Fig S1C-F**).

### LRP1 deficiency in skeletal progenitors results in dwarfism, impaired mobility, abnormal gaits and fore/hind limb malformation

We next investigated the impact of early skeletal developmental defects in postnatal *Lrp1*^flox/flox^*/Prrx1*^Cre^ mice. The mice survived into adulthood, but the weight of the homozygote mice was consistently lower than their WT and heterozygote littermates starting 3 weeks up to 14 weeks after birth (**Fig 3A**). Strikingly, *Lrp1*^flox/flox^*/Prrx1*^Cre^ mice consistently showed altered posture and impaired mobility at birth which was persistent up until 14 weeks of age (**Movies S1 and S2**). A continuous automated home-cage monitoring system for a 4-week period from week 6 showed significantly reduced locomotor activity of *Lrp1*^flox/flox^*/Prrx1*^Cre^ mice compared with WT littermates (**Fig 3B**). Whole-mount skeletal staining of 14-week-old postnatal adult mice with Alcian blue and Alizarin red showed that *Lrp1*^flox/flox^*/Prrx1*^Cre^ mice had shorter limbs and smaller stature than their WT littermates (**Fig 3C**). *Lrp1*^flox/flox^*/Prrx1*^Cre^ mice were unable to open digits in their hind limb (**Fig 3D**). As an occasional feature in *Lrp1*^flox/flox^*/Prrx1*^Cre^ mice, fused and abnormal digits were observed in their fore limb (**Fig 3EF**). None of these phenotypes were observed in *Lrp1*^flox/flox^*/Prrx1*^Cre^ heterozygote mice, suggesting an autosomal recessive phenotype.

**Fig 3.**
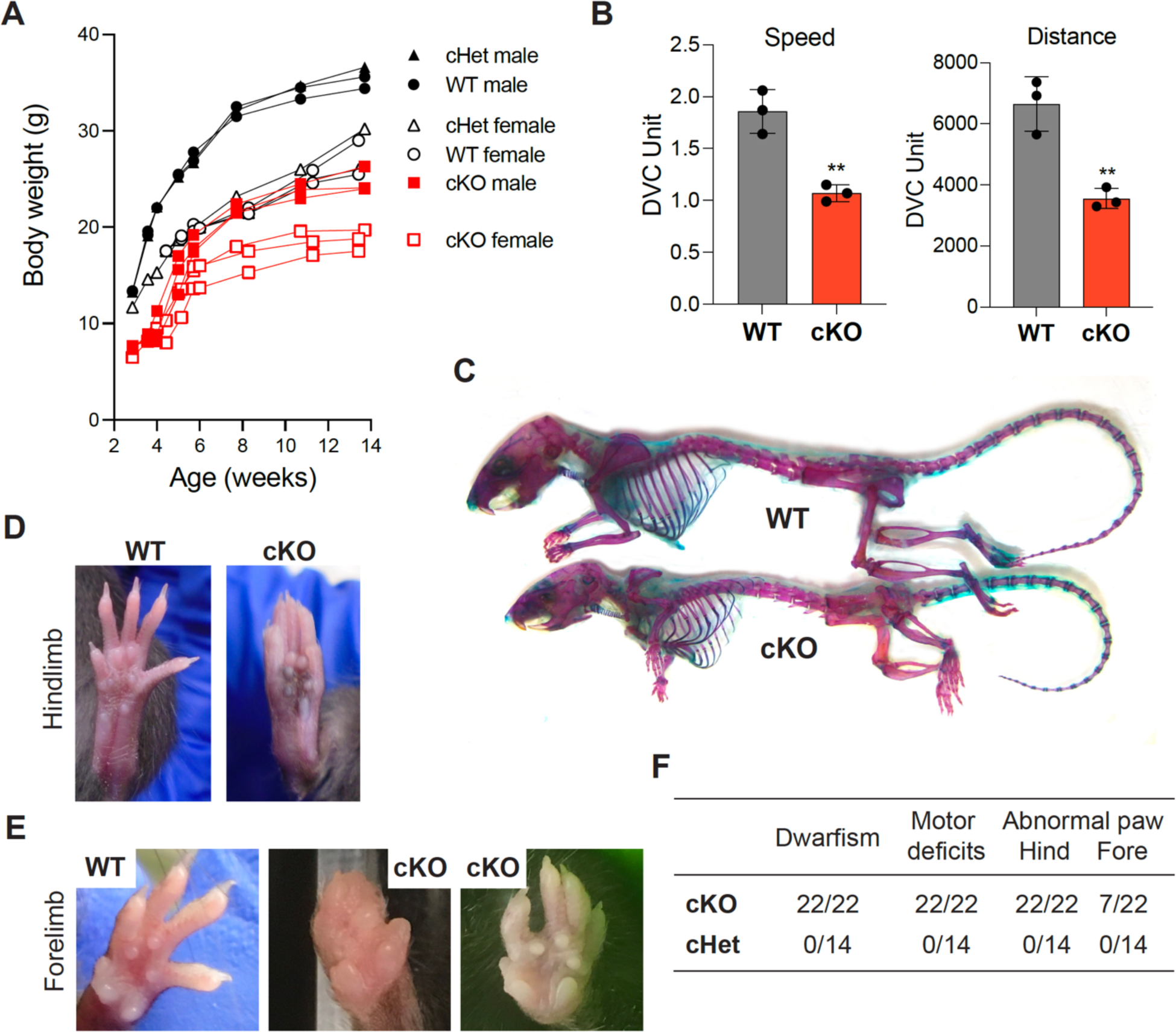
LRP1 deficiency in skeletal progenitors results in dwarfism, impaired mobility and fore/hind limb malformation. *A*, Mouse weight measurement from 3-14 weeks after birth. Closed (male) and open (female) black circles (WT), black triangles (cHet; *Lrp1*^flox/flox^*/Prrx1*^Cre^ heterozygote conditional KO), and red squares (cKO; *Lrp1*^flox/flox^*/Prrx1*^Cre^ homozygote conditional KO) represent individual mice. *B*, Total distance and average speed of WT and cKO mice. Locomotor activity of 6-week-old mice measured by the continuous automated home-cage monitoring system for 4 weeks. Circles represent individual mice and bars show the mean ± *SD*. **, *p* < 0.01 by 2-tailed Student’s t test. *C*, Whole-mount skeletal staining of 14-week-old mice with Alcian blue and Alizarin red. *D*, Photographs of 8-week-old WT and the cKO hind paws. *E*, Photographs of 8-14-week-old WT and the cKO fore paws. *F*, Table showing the frequencies of the identified phenotypes.

### Severe and persistent defects in multiple bones and joints in *Lrp1*^flox/flox^*/Prrx1*^Cre^ mice

*In vivo* µCT scanning of 2-week-old *Lrp1*^flox/flox^*/Prrx1*^Cre^ mice revealed that long bones were not only shorter but markedly thicker and twisted, with much delayed or a lock of emergence of secondary ossification centres (**Fig 4A**). The upper limbs were abnormally twisted and the scapular were poorly defined compared with WT. The absence of crescent-shaped acetabulum sockets, rounded femoral heads and patella bone in *Lrp1*^flox/flox^*/Prrx1*^Cre^ mice suggested an impaired mobility. Notably, these bone and joint defects remained defective in 14-week-old mice (**Fig 4B**). The observed difference in the length and width of femur and humerus bones between 6-week-old WT and *Lrp1*^flox/flox^*/Prrx1*^Cre^ mice persisted in 14-week-old mice (**Fig 4C** and **S2**). High-resolution µCT scanning of a 14-week-old *Lrp1*^flox/flox^*/Prrx1*^Cre^ tibia revealed a substantial reduction in trabecular bone density with virtual absence of primary spongiosa compared to the WT tibia (**Fig 4D**). Fusion and deformities of phalanges were observed in *Lrp1*^flox/flox^*/Prrx1*^Cre^ forelimb with fused and abnormal digits (**Fig 4E**).

**Fig 4.**
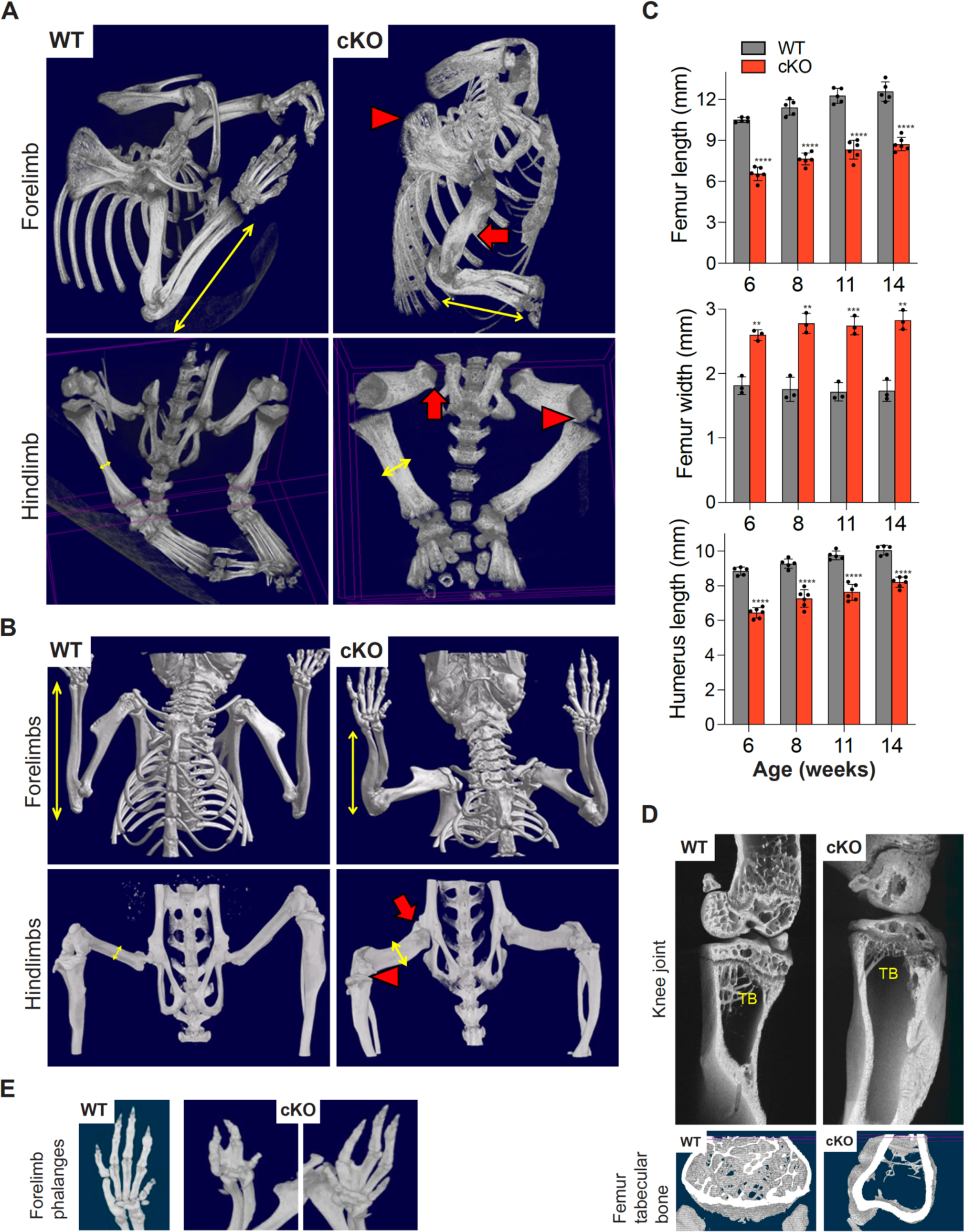
Severe and persistent defects in multiple bones and joint in *Lrp1*^flox/flox^*/Prrx1*^Cre^ mice. *A and B*, Representative images of in vivo X-ray analysis of limbs of 2-week-old (*A*) and 14-week-old (*B*) WT and *Lrp1*^flox/flox^*/Prrx1*^Cre^ homozygote conditional KO (cKO) mice. Yellow arrows indicate difference of bone length (top panels) and width (bottom panels) between WT and cKO mice. Red arrowheads and arrows indicate undefined shoulder blades and twisted long bones (top panels), and undefined knee and lack of hip joints (bottom panels) in cKO mice, respectively. *C*, Femur bone length and width of 6-14-week-old mice. Circles represent individual mice and bars show the mean ± *SD*. **, *p* < 0.01; ***, *p* < 0.001; ****, *p* < 0.0001 by 2-tailed Student’s t test. *D*, Representative images of high-resolution μCT analysis of femur trabecular bones of 14-week-old mice. TB, trabecular bone. *E*, Representative images of in vivo X-ray analysis of forelimb phalanges of 14-week-old cKO mice.

### Defects in growth plate, organisation of columnar chondrocytes, secondary ossification, articulation and cavitation of joints and proteoglycan turnover in *Lrp1*^flox/flox^*/Prrx1*^Cre^ mice

To investigate details of the multiple and severe skeletal defects in *Lrp1*^flox/flox^*/Prrx1*^Cre^ mice, we histologically examined the bone and joint sections. H&E staining of 2- and/or 14-week-old *Lrp1*^flox/flox^*/Prrx1*^Cre^ shoulder, elbow and knee revealed striking defects (**Fig 5A-C**). These included a deformed and disrupted growth plate with poorly stacked columnar chondrocytes, markedly delayed secondary ossification, impaired articulation as a result of fused bone ends and lack of articular cartilage formation. Safranin-O/fast green staining further revealed proteoglycan depletion in articular cartilage and the growth plate in adulthood at 14-week-old mice but not during growth phase in 2-week-old mice (**Fig 5A-C**). Similar results were also obtained by Toluidine blue staining. Notably, femoral heads were substantially deformed with an extra groove juxtaposed to a poorly developed acetabulum socket (**Fig 5D**). Sox9, a pivotal transcription factor expressed in developing and adult cartilage (40–42), was almost absent in the articular cartilage and growth plate of 14-week-old *Lrp1*^flox/flox^*/Prrx1*^Cre^ compared with WT mice (**Fig 5E**), emphasising defects in cartilage development.

**Fig 5.**
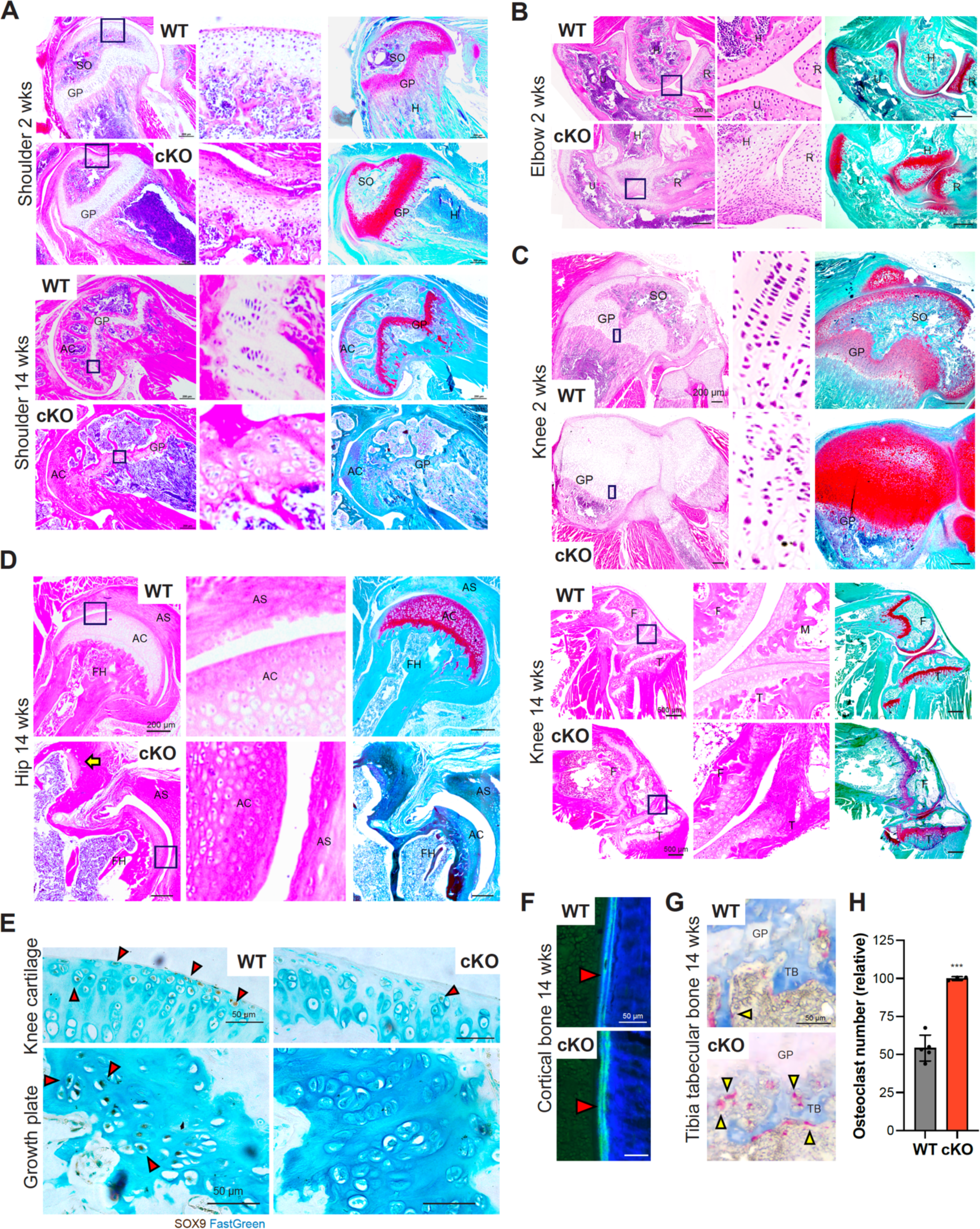
Defects in growth plate, organisation of columnar chondrocytes, secondary ossification, articulation and cavitation of joints and proteoglycan turnover in *Lrp1*^flox/flox^*/Prrx1*^Cre^ mice. *A*-*D*, Representative images of H&E and safranin-O/fast green staining in shoulder (*A*), elbow (*B*), knee (*C*) and hip (*D*) joint sections of 2- and/or 14-week-old WT and *Lrp1*^flox/flox^*/Prrx1*^Cre^ (cKO) mice. Arrow indicates the extra groove of femoral heads (*D*). Regions delineated by the dark blue squares in the panels have been magnified in the right panels. Scale bar, 200 µm. GP, growth plate; AC, articular cartilage; F, femur; T, tibia; M, menisci. *E*, Representative images of immunohistochemical staining of SOX9 in knee articular cartilage and growth plate sections of 14-week-old WT and cKO mice. Arrow heads indicate cells with SOX9 positive staining. Scale bar, 50 µm. *F*, Representative images of fluorescent microscopy analysis of calcein-double stained cortical bones in 14-week-old mice (n = 3 each). Red arrowheads indicate calcein double staining in cortical bone. Scale bar, 50 µm. *G*, TRAP and aniline blue staining of tibia trabecular bone of 14-week-old mice. Yellow arrowheads highlight osteoclast staining. Scale bar, 50 µm. TB, trabecular bone; GP, growth plate. *H*, Measurement of osteoclast numbers in WT and *Lrp1*^flox/flox^*/Prrx1*^Cre^ (cKO) trabecular bones. Circles represent individual mice and bars show the mean ± *SD*. ***, *p* < 0.001 by 2-tailed Student’s t test.

We then tested dynamic histomorphometry of bone formation using calcein-double labelling in 14-week-old mice and showed a comparable rate of bone formation rate between WT and *Lrp1*^flox/flox^*/Prrx1*^Cre^ mice (**Fig 5F** and **S3A-C**). Further analysis using tartrate-resistant acid phosphatase (TRAP) staining for osteoclasts revealed approximately 1.9-fold increased osteoclast activity in 14-week-old *Lrp1*^flox/flox^*/Prrx1*^Cre^ compared with WT littermates (**Fig 5GH** and **S3DE**). These results were consistent with *Lrp1* deficiency in osteoblasts (28) or osteoclasts (29) resulting in low bone-mass phenotype due to increased osteoclastogenesis.

### LRP1 mediates endocytic recycling of Wnt5a, a core non-canonical WNT/planar cell polarity (PCP) pathway component

The observed skeletal defects could potentially arise from a range of mechanisms. Markedly thicker and shorter long bones (**Fig 4**), and disrupted growth plate with disorganised columnar chondrocytes (**Fig 5**) closely resemble phenotypes associated with defects in the non-canonical WNT/PCP signalling pathway (**Fig 6A**)(43). For example, deletion of core components of WNT/PCP including Ror2 (44), Vangl2 (45), Ryk (46) and Wnt5a (47) as well as inducible Wnt5a overexpression (48) all resulted in shorter and thicker long bones. Our recent study identified >50 novel candidate binding partners for LRP1 in cartilage (9) including Wnt5a and Wnt11, key components of WNT/PCP signalling. We therefore investigated the potential for direct interaction of Wnt5a and Wnt11 with LRP1 using a solid-phase binding assay. The assay employed purified Wnts and full-length LRP1(49). We found that Wnt5a and Wnt11 directly bind to immobilised LRP1 with high affinity (apparent binding constant (*K_D,app_*) of 31 nM and 42 nM, respectively (**Fig 6BC**). In contrast, binding of Wnt3a, a key canonical WNT component, to LRP1 was negligible with *K_D,app_* of >200 nM (**Fig 6D**).

**Fig 6.**
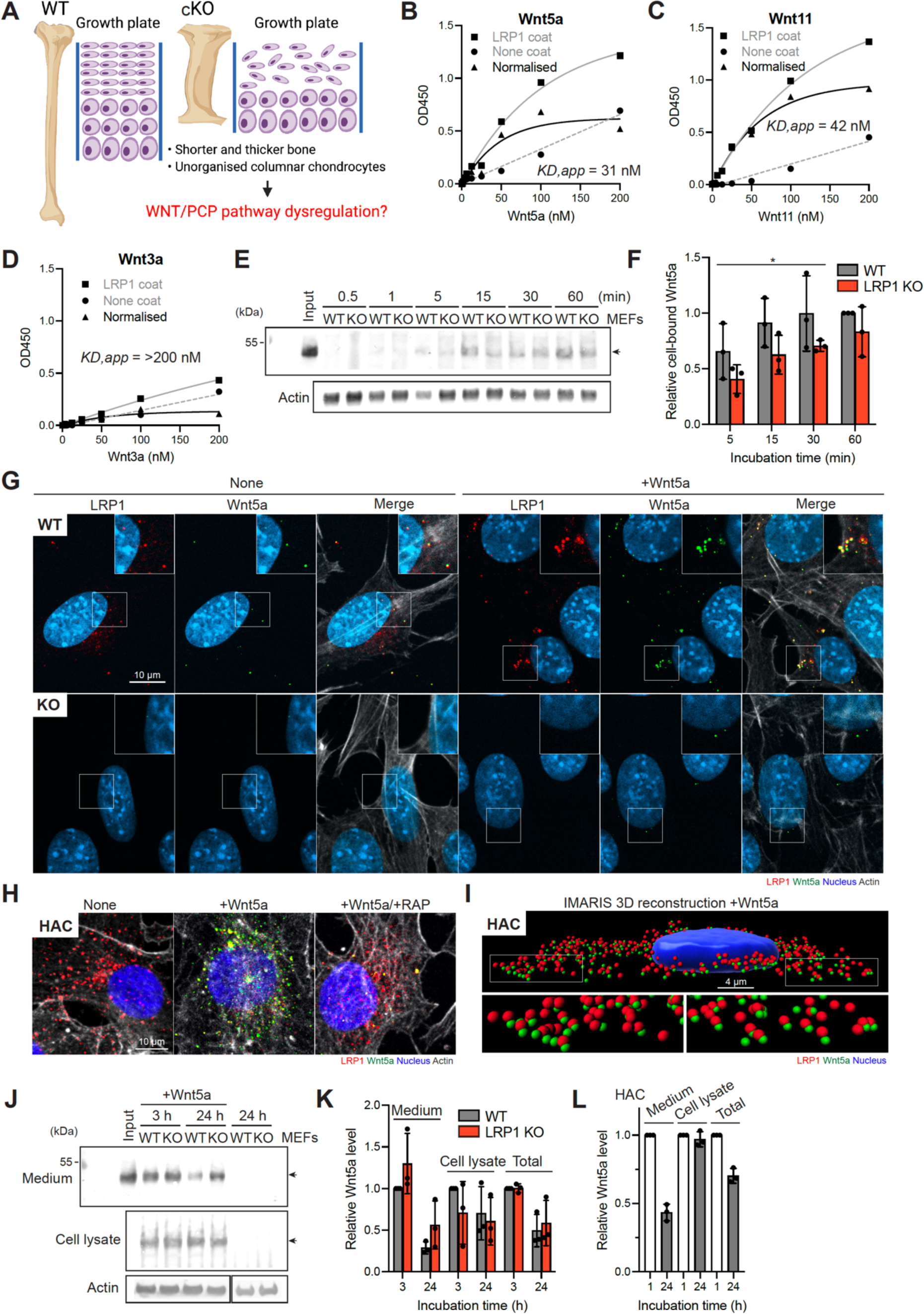
LRP1 mediates endocytic recycling of Wnt5a, a core non-canonical WNT/planar cell polarity (PCP) pathway component. *A*, Schematic diagram illustrating the long bone phenotype of *Lrp1^flox/flox^/Prrx1^Cre^* mice. *B-D*, Full-length sLRP1 was coated onto microtiter plates and the binding of 0-200 nM Wnt5a (*B*), Wnt11 (*C*), Wnt3a (*D*) was measured using specific antibody for each Wnt as described under “Materials and Methods”. Mean values of technical duplicates for none-coating, LRP1-coating and after normalisation were shown as circles, squares and triangles, respectively. Extrapolated *K_D,app_* values were estimated based on one-phase decay nonlinear fit analysis (black lines). *E and F*, WT and LRP1 KO MEFs (n = 3) were incubated with 40 nM Wnt5a for 0.5-60 min and Wnt5a in the cell lysate was detected by Western blotting (*E*). The relative amount of Wnt5a was expressed by taking the amount of Wnt5a after 60-min incubation as 1 (*F*). Circles represent individual mice and bars show the mean ± *SD*. *p* values for the amount of Wnt5a after incubation for 5-30 min in WT *versus* LRP1 KO MEFs were evaluated by two-way ANOVA. *, *p* < 0.05. *G and H*, Representative images of confocal microscopy analysis for Wnt5a and LRP1 in WT and LRP1 KO MEFs (n = 3) (*E*) or human normal chondrocytes (n = 3) (*F*). Cells were incubated with 20 nM Wnt5a for 3 h in the absence (*G and H*) or presence of 500 nM RAP (*H*). Wnt5a, LRP1, cytoskeleton and nucleus were visualised as described under “Materials and Methods”. Scale bar, 10 µm. *I*, Imaris image visualisation of staining of LRP1 and Wnt5a in human chondrocyte incubated with 20 nM Wnt5a for 3 h. Scale bar, 4 µm. Regions delineated by the white squares in the panels have been magnified in the top right (*G*) or the bottom (*I*). *J and K*, WT and LRP1 KO MEFs (n = 3) were incubated with 20 nM Wnt5a for 3-24 h and Wnt5a in the medium and cell lysate were detected by Western blotting (*J*). Densitometric analysis of immunoreactive Wnt5a bands was carried out. The relative amount of Wnt5a in the media, cell lysate and both media and cell lysate (total) were expressed by taking the amount of Wn5a after 3-h incubation in WT MEFs as 1 (*K*). *L*, human normal chondrocytes (n = 3) were incubated with 20 nM Wnt5a for 1-24 h and analysed as in *A* and *B*. The relative amount of Wnt5a after 24-h incubation was expressed by taking the amount of Wn5a after 1-h incubation as 1. Circles represent individual experiment and bars show the mean ± *SD*.

Since Wnt5a dysregulation may explain the malformation of long bones, function of LRP1 in cellular trafficking of Wnt5a was investigated. To test whether LRP1 facilitates cell-association of Wnt5a through direct capture of Wnt5a, exogenously added Wnt5a in the cell-lysate of WT and LRP1 KO mouse embryonic fibroblasts (MEFs) (50) was monitored for incubation periods ranging between 0-60 min. The levels of cell-bound Wnt5a in WT MEFs after incubation for 5, 15 and 30 min were significantly higher than these in LRP1 KO cells with 1.6, 1.5 and 1.4-fold differences, respectively (**Fig 6EF**). We next examined internalisation of Wnt5a in these cells. Immunofluorescent confocal microscopy analysis revealed the co-localisation of Wnt5a and LRP1 inside WT MEFs (**Fig 6G**). The fluorescent signal of Wnt5a was markedly reduced in the LRP1 KO MEFs, indicating that LRP1 is responsible for cellular update of Wnt5a. The intracellular colocalisation of exogenously added Wnt5a and LRP1 was also observed in human primary chondrocytes (**Fig 6H**). This was markedly reduced in the presence of receptor-associated protein (RAP), which antagonises ligand binding by competitively occupying the ligand binding region of LRP1. Three-dimensional reconstruction of the confocal microscopy images using IMARIS (v8.1) further confirmed that the majority of internalised Wnt5a was associated with LRP1 (**Fig 6I**).

Several secreted LRP1 ligands are degraded intracellularly following internalisation (6–10). Some ligands, however, are recycled back to the extracellular milieu, facilitating their distribution and availability (51, 52). As mentioned above, Wnt5a was identified as LRP1 ligand candidate in our previous study based on that it was co-immunoprecipitated from the chondrocyte culture medium with recombinant soluble LRP1 ligand-binding cluster II (sLRP1-II) (9), which is considered to be responsible for most ligand binding (7, 8, 53–56). However, our secretome analysis found that Wnt5a is not increased in the chondrocyte culture medium upon inhibition of LRP1-mediated endocytosis (9), suggesting a possibility of its endocytic recycling. We thus monitored the level of exogenously added Wnt5a in the medium and cell lysate of WT and LRP1 KO MEFs by Western blot analysis. After 3-24 h incubation, exogenously added Wnt5a was detected in both the conditioned medium and cell lysate (**Fig 6JK**). No significant difference in Wnt5a levels were observed in LRP1 KO compared to WT MEFs and relatively large amounts of Wnt5a still remained even after 24-h incubation with the cells. Human chondrocytes express higher levels of LRP1 compared to WT MEFs but unlike other LRP1 ligands such as TIMP3 (5, 6), MMP13 (7) and a disintegrin and metalloproteinase with thrombospondin motifs (ADAMTS)5 (10), rapid clearance of Wnt5 was not observed in human chondrocytes (**Fig 6L**). These results suggest that LRP1 facilitates cell-association and mediates internalisation of Wnt5a but not its intracellular degradation.

### LRP1 regulates WNT/PCP signalling in *Xenopus* embryonic development

To evaluate the significance of LRP1-Wnt5a interaction in the WNT/PCP signalling pathway, we employed *Xenopus laevis* (African clawed frog), which is an invaluable model system for studying the role of WNT signalling in development (57). WNT/PCP singnalling directly controls convergent extension movements in the developing embryo. Loss of expression or overexpression of WNT/PCP components leads to embryos showing shortened trunks (**Fig S4A**)(58, 59). We confirmed that injection of Wnt5a or Wnt11 mRNAs into the dorsal marginal zone of 4-cell stage embryos results in shortened tadpoles (**Fig S4BC**). According to Xenbase, *lrp1* gene expression in Xenopus laevis starts at Oocyte V-VI stage, peaks at Nieuwkoop and Faber stage 1 and then remains on throughout development (**Fig 7A**)(60, 61). Wholemount *in situ* hybridization using a *lrp1* probe showed that *lrp1* is expressed in the neural tube, branchial arches, somites and neuroadrenergic cells during the development (**Fig 7B**). Gene-knockdown of *lrp1* by injection of various doses of lrp1 morpholino into the dorsal side of 4-cell stage embryos caused conversion extension phenotypes of shortened trunks at all the concentrations tested (**Fig 7CD)**. Compared to control morpholino, 20 ng of lrp1 morpholino increased frequency of conversion extension phenotype by ∼2.2-fold (**Fig 7E)**. We next examined the effect of overexpression of LRP1 mini-receptor consisting of the ligand-binding cluster II and the entire C-terminus, including the transmembrane domain and the cytoplasmic tail. This functional mini-LRP1 receptor maintains capacity to mediate clathrin-dependent endocytosis of LRP1 ligands (62). Injection of the mini-LRP1 increased frequency of conversion extension phenotype in a dose-dependent manner (**Fig 7F**). Compared to control, 5 pg of mini-*Lrp1* injection increased frequency of conversion extension phenotype by ∼8.2-fold (**Fig 7G)**. These results suggest a role for LRP1 in regulation of the WNT/PCP pathway.

**Fig 7.**
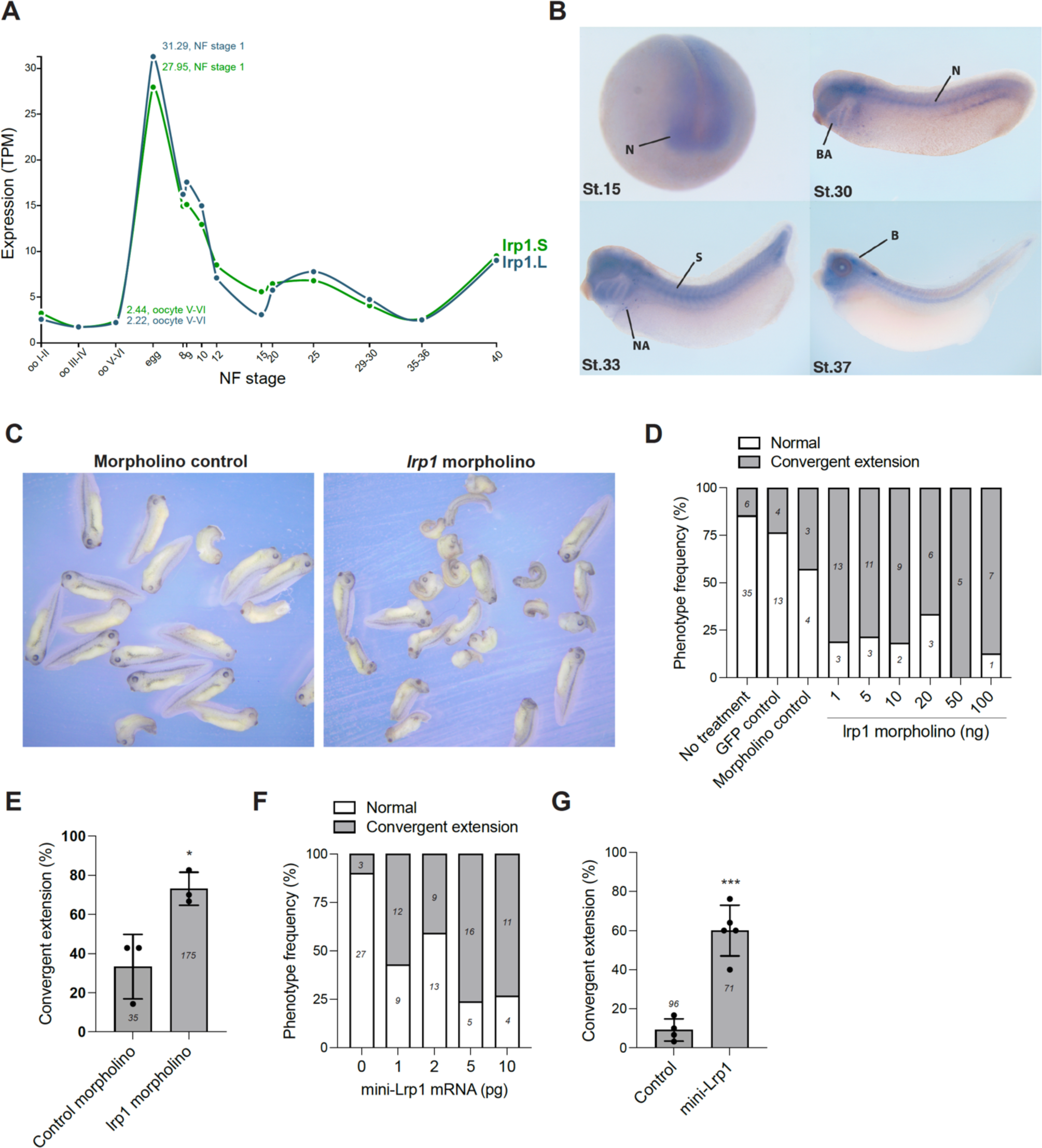
LRP1 regulates WNT/PCP signalling in *Xenopus* embryonic development. *A*, *lrp1* gene expression profile during *Xenopus laevis* embryonic development available in Xenbase. NF, Nieuwkoop and Faber. *B*, *lrp1* wholemount in situ hybridisation during the development. N, neural tissue; BA, branchial arches; S, somites; NA, neuroadrenergic cells; B, brain. *C-E*, Various doses (*D*) or 20 ng (*C* and *E*) of *lrp1* morpholino were injected into 1 cell of 4-cell stage embryos in the dorsal marginal zone. Representative images of Xenopus tadpoles treated with control and *lrp1* morpholino after fixation at NF stage 35/36 (*C*). Normal embryos and those with convergent extension phenotype were counted and percentages for frequency of each phenotype (*D*) or convergent extension phenotype in three independent experiments (*E*) were shown. Circles represent individual experiment and bars show the mean ± *SD*. *, *p* < 0.05 by 2-tailed Student’s t test. Number of embryos counted is stated in each bar graph. *F* and *G*, Various doses (*F*) or 5 pg (*G*) of mini-*Lrp1* mRNA were injected and the phenotypes were counted as in *D* and *E*. Circles represent individual experiment and bars show the mean ± *SD*. ***, *p* < 0.001 by 2-tailed Student’s t test. Number of embryos counted is stated in each bar graph.

### LRP1 partially colocalises with Wnt5a and its deficiency diminishes graded distribution of Wnt5a and Vangl2 in the developing limbs

Lastly, we investigated the interaction of LRP1 and Wnt5a and how LRP1 deficiency affects Wnt5a distribution and activity in the developing limbs. Wnt5a mRNA is expressed in the perichondrium in developing limbs (47, 63, 64) but Wnt5a protein distribution in the developing limbs remains incompletely understood. Our histological investigation of Wnt5a protein in E16.5 elbow showed a Wnt5a distribution pattern similar to that seen for LRP1; abundant expression of Wnt5a in perichondrium and proliferative flattened chondrocytes, but very weak expression in hypertrophic chondrocytes (**Fig 8A**). Immunofluorescent confocal microscopy analysis further revealed that partial colocalisation of Wnt5a and LRP1 in E16.5 hind limbs (**Fig 8B**). Notably, Wnt5a immunosignal was substantially reduced in E16.5 *Lrp1*^flox/flox^*/Prrx1*^Cre^ compared to WT limbs. To evaluate Wnt5a activity, we examined expression and phosphorylation of Vangl2, a core component of WNT/PCP as mentioned above. It has been reported that the abundance of Vangl2 is tightly controlled by the ubiquitin-proteasome system through endoplasmic reticulum–associated degradation (65). Wnt5a activity prevents proteasomal degradation of Vangl2, facilitating its export from the endoplasmic reticulum to the plasma membrane. Wnt5a also dose-dependently induces phosphorylation of Vangl2 through Ror2 to establish WNT/PCP gradient (45). Immunofluorescent confocal microscopy analysis revealed a remarkable reduction of Vangl2 in E16.5 *Lrp1*^flox/flox^*/Prrx1*^Cre^ compared to WT limbs (**Fig 8C**). Immunosignal of phosphorylated Vangl2 was detected in E16.5 WT elbow, whereas it was almost absent in *Lrp1*^flox/flox^*/Prrx1*^Cre^ limbs (**Fig 8D**). These results suggest that LRP1 deficiency causes aberrant WNT/PCP signalling.

**Fig 8.**
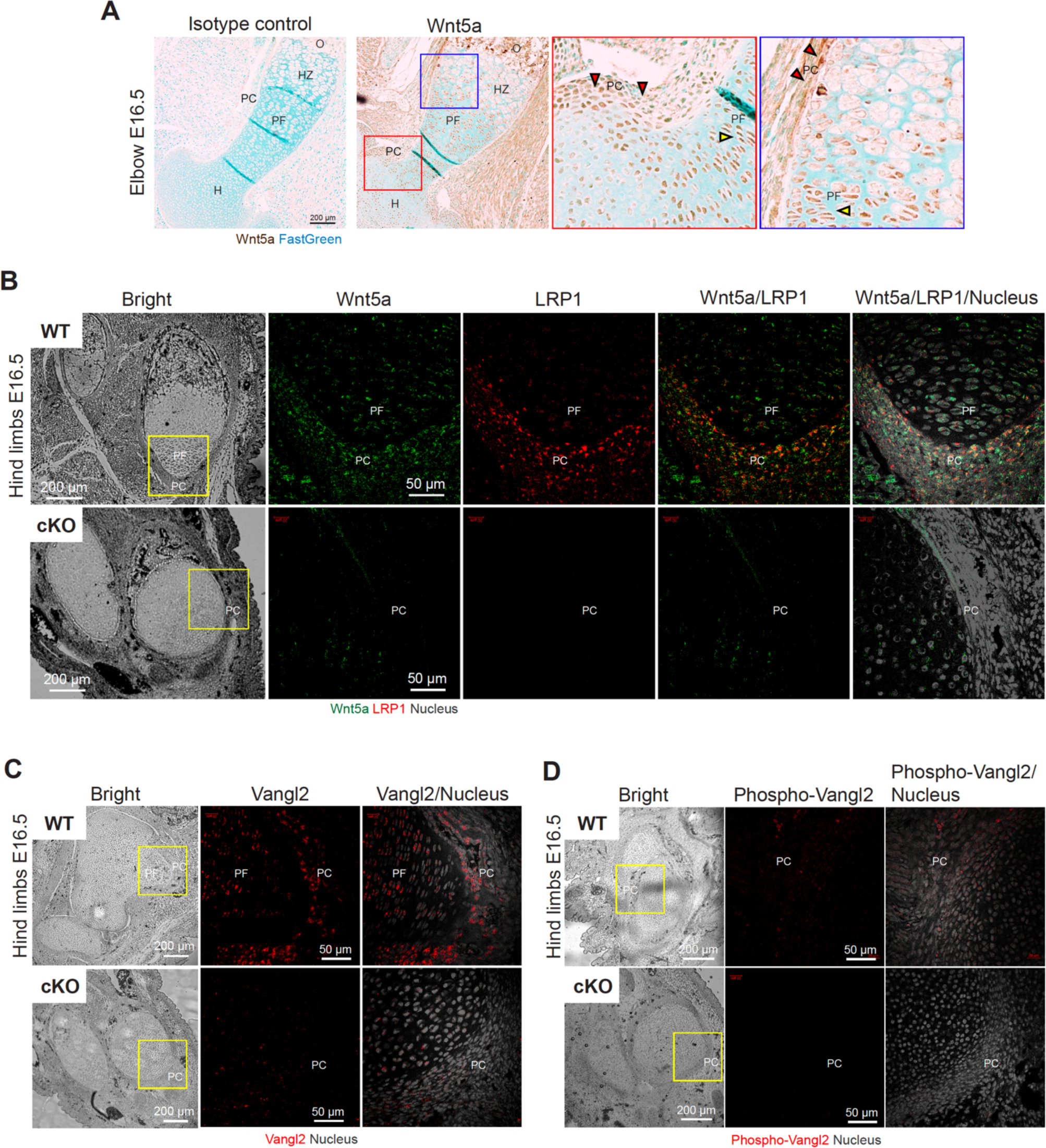
LRP1 partially colocalises with Wnt5a and its deficiency diminishes graded distribution of Wnt5a and Vangl2 in the developing limbs. *A*, Representative images of immunohistochemical staining of Wnt5a and fast green counterstaining in E16.5 elbow sections of WT (n = 3). Scale bar, 200 µm. Regions delineated by the squares have been magnified in the rght panels. Red and tallow arrowheads indicate Wnt5a protein in perichondrium and proliferative flattened chondrocytes, respectively. H, humerus; HZ, hypertrophic zone; O, primary ossification center; PF, proliferative flattened chondrocytes; PC, perichondrium. *B-D*, Representative images of confocal microscopy analysis for Wnt5a and LRP1 (*B*), Vangl2 (*C*) and phosphorylated Vangl2 (*D*) in E16.5 hind limb sections of WT and *Lrp1*^flox/flox^*/Prrx1*^Cre^ (cKO) mice (n = 3). Wnt5a, LRP1, Vangl2, phospho-Vangl2 and nucleus were visualised as described under “Materials and Methods”. Regions delineated by the yellow squares in the bright field images have been magnified in the left panes. PF, proliferative flattened chondrocytes; PC, perichondrium. Scale bar, 50 µm.

## Discussion

This study showed, for the first time, a critical role of LRP1 in skeletal progenitor cells and its regulation of the WNT/PCP pathway. *Lrp1* deletion in early skeletal progenitors caused severe defects in multiple bones and joints which persisted into skeletally mature 14-week-old mice, indicating a non-redundant function in skeletal development and maturity. These observations of long bone and joint malformations were not evident in *Lrp1*^flox/flox^*/Col2a1*^Cre^ mice (27) or *Lrp1*^flox/flox^/*Acan*^CreERT2^ mice. In the *Lrp1*^flox/flox^*/Col2a1*^Cre^ strain, *Col2a1*^Cre^ gene expression was confirmed in limbs at as early as E12.5 (66), whereas in our *Lrp1*^flox/flox^*/Prrx1*^Cre^ mice, *Prrx1*^Cre^ gene expression starts at E9.5 (35–38). These studies suggest that LRP1 in early skeletal progenitor cells prior to E12.5 is more intrinsically involved in guiding proper mesenchymal cell recruitment for the formation of multiple cartilage and bone elements. Similarly, mice, in which Wnt5a overexpression is induced at E10.5 exhibited a much more severe long bone phenotype than mice induced at E12.5 (48). Indeed, overexpression of Wnt5a at E13.5 does not result in visible bone phenotype, indicating that the most critical period for Wnt5a in limb development is prior to E13.5. Our study using different models demonstrate the regulation of WNT/PCP signaling pathway by LRP1 during this critical early time point.

Skeletal progenitor LRP1 has at least two different functions: to remove molecules from the extracellular milieu and target them for intracellular degradation, and to capture, recycle and distribute molecules (**Fig 9**). LRP1 phosphorylation is likely to be dispensable for its function in skeletal development since knock-in mouse models of mutation of the proximal, and separately, distal NPxY motif to disable phosphorylation exhibited normal skeletal development (67). Our data clearly demonstrate that TIMP3 and CCN2 are tightly regulated LRP1 ligands (6, 9), which accumulated throughout the limb in P0 when LRP1 was deleted in *Lrp1*^flox/flox^/*Prrx1*^Cre^ mice. This indicated that in addition to the WNT/PCP pathway regulation, LRP1 is important in regulating extracellular concentrations of biologically active ligands such as TIMP3 and CCN2 in the developing limbs.

**Fig 9.**
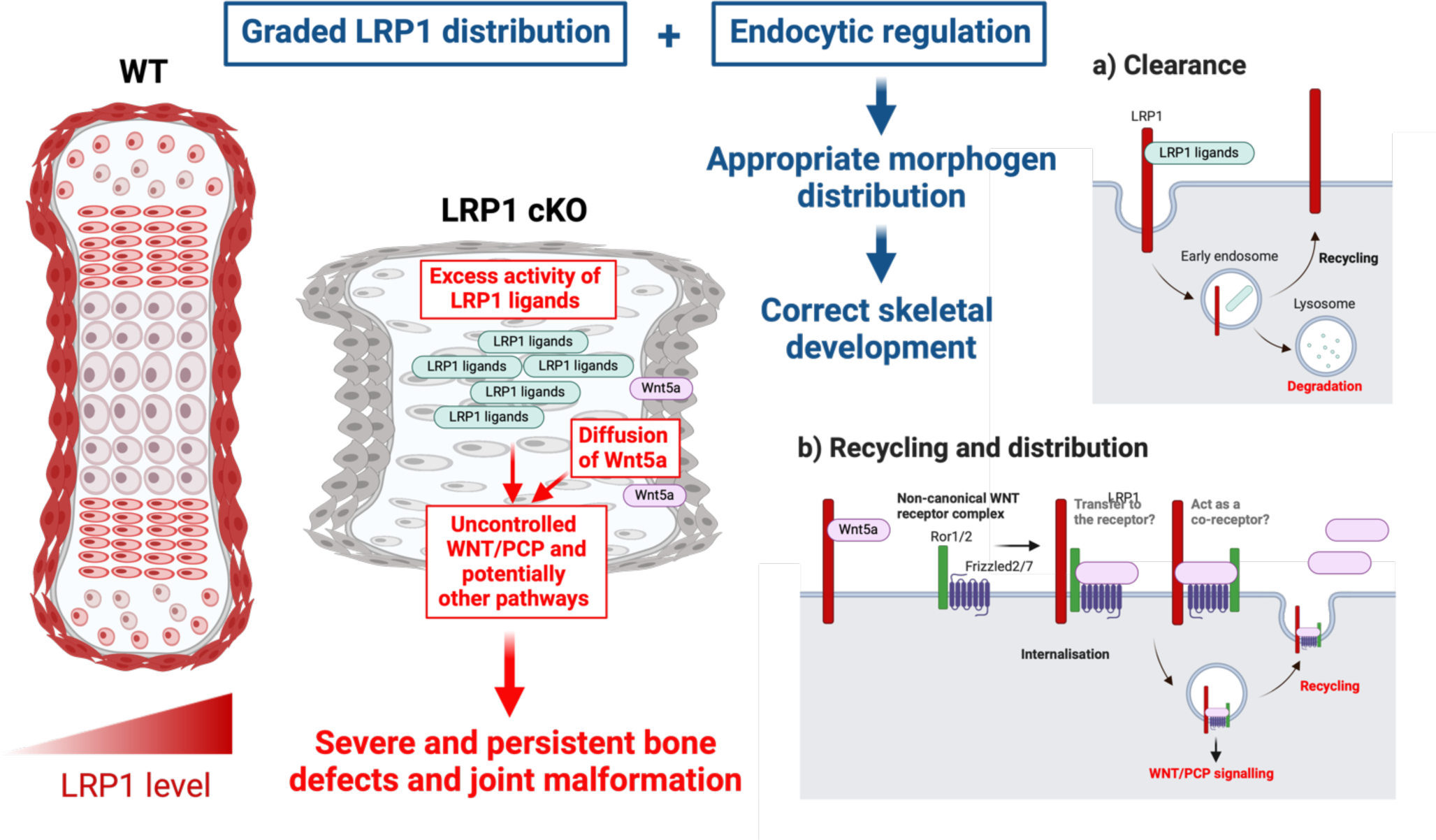
A novel and critical role for LRP1 in skeletal development and its deficiency in emergence of skeletal pathologies. Skeletal progenitor LRP1 is likely to have at least two different functions: to remove molecules from extracellular milieu and degrad them intracellularly (a), and to capture, recycle and distribute molecules (b). Combination of graded distribution of LRP1 and LRP1-mediated endocytosis regulates distribution of extracellular signalling molecules. This provides a novel mechanism for appropriate distribution of extracellular signalling molecules to ensure that bone and joint form correctly. Thus, loss of LRP1 leads to excess activity of some of LRP1 ligands and erratic concentrations of Wnt5a, resulting in multiple and severe bone and joint defects.

This study identifies LRP1 as a major receptor for Wnt5a. Higher levels of cell-associated Wnt5a were detected in WT compared with LRP1 KO cells, suggesting that LRP1 effectively captures Wnt5a from the cellular microenvironment. It has been reported that Wnt5a is internalised by cells *via* a clathrin-dependent pathway (68, 69). We found that LRP1 mediates Wnt5a endocytosis but internalised Wnt5a is not degraded, and is most likely recycled. Our immunofluorescence staining further revealed a colocalisation of LRP1 and Wnt5a in bone template and perichondrium of E16.5 WT limbs. Strikingly, Wnt5a and Vangl2 was almost diminished in E16.5 *Lrp1*^flox/flox^/*Prrx1*^Cre^ limbs, suggesting that LRP1 loss causes dysregulation of the WNT/PCP pathway. We are currently exploring two possibilities. One is that LRP1 facilitates Wnt5a binding to its cell-surface receptor complexes consisting of Ror1/2 and the frizzled receptors (**Fig 9B**)(70–72). The other is that LRP1 directly interacts with and stabilises Vangl2.

The perichondrium is a dense layer of fibrous connective tissue that covers cartilage in endochondral ossification. Two-way signalling between cells in the perichondrium and the underlying cartilage are essential for endochondral bone formation but questions remain regarding molecular mechanisms underpinning cellular communication between them (34, 43). Our present study revealed an abundant expression of LRP1 in the perichondrium, strongly suggesting a role for LRP1 in the signaling pathways underpinning bond formation. Furthermore, the colocalization of LRP1 and Wnt5a in the perichondrium raises the possibility that these two molecules interact to control the recruitment of chondroprogenitor cells from the perichondrium into growth plate. We hypothesise that absence of their interaction in *Lrp1*^flox/flox^*/Prrx1*^Cre^ limbs results in uncontrolled recruitment of chondroprogenitor cells leading to skeletal element widening.

It is worth noting that the data presented here differ significantly from other LRP family members associated with skeletal formation, such as LRP5 and LRP6, which function as Wnt coreceptors with the frizzled receptor and regulate canonical WNT/β-catenin signalling (73, 74). LRP5/6 interact with many Wnts (Wnt1/2/3/3a/2b/6/8a/9a/9b/10b) but not Wnt5a or Wnt11 (75). *Lrp6* KO mice die at birth and exhibit a variety of severe developmental abnormalities resembling those caused by mutations in Wnt1, Wnt3a and Wnt7a (73). *Lrp6* KO limbs showed a consistent loss of the most posterior digit but did not exhibit the limb outgrowth defects observed in *Wnt5a* KO mice (47). Some *Lrp6* KO mice exhibited deletion of additional digits and the radius, as well as malformation of the ulna. Therefore, *Lrp1*^flox/flox^*/Prrx1*^Cre^ mice *Lrp6* KO phenotypes are distinct.

In *Xenopus*, convergent extension movements are known to be controlled by non-canonical Wnt signalling (58, 59). We clearly show that *lrp1* knockdown using morpholinos or overexpression by using a well characterised mini-Lrp1 construct leads to convergent extension phenotypes similar to Wnt5a and Wnt11 (**Fig 7 C-G** and **S4**). These results suggest a clear interaction of lrp1 with non-canonical Wnt signalling. Nonetheless, accumulating evidence suggests a role of LRP1 in the regulation of canonical WNT signalling. *Lrp1* deletion in mouse neural crest cells results in heart defects, which are associated with a decrease in canonical Wnt signalling (22). In vitro, LRP1 interacts with Frizzled1 and downregulates WNT/β-catenin pathway in HEK293T cells (76). In contrast, LRP1 stimulates WNT/β-catenin pathway and prevents intracellular cholesterol accumulation in fibroblasts (77). Macrophage LRP1 also increases WNT/β-catenin pathway by directly binding to and effectively removing secreted frizzled-related protein 5, which prevents Wnt binding to its receptor (78). These studies emphasise the developmental stage and tissue-specific nature of WNT signalling. Considering the importance of WNT/β-catenin pathway for synovial joint formation (79), LRP1 may selectively regulate a canonical WNT singalling during synovial joint formation, which requires further investigations.

Since the 2.4 kb *Prrx1* enhancer is reported to be expressed in skeletal muscle as early as E16.5 (37), we cannot exclude the possibility that skeletal muscle LRP1 plays a role in the observed phenotype. In particular twisted long bones, can be explained by altered mechanical adaptation due to abnormal skeletal muscle-bone interaction. *Lrp1*^flox/flox^/*Prrx1*^Cre^ mice exhibited significant differences in limb length and femur thickness as early as at P0. One possibility to rationalise drastic skeletal changes at this stage is the mechanical stimuli generated by the embryo’s active movements. Embryos with restricted movements in utero showed a skeletal phenotype of reduced ossification, especially in forelimbs (80).

In conclusion, we showed that abundant expression of LRP1 in early skeletal precursor cells, its critical role for synovial joint formation and accurate bone growth. We further demonstrated the regulation of WNT/PCP signaling pathway by LRP1 which may explain the malformation of long bones in *Lrp1*^flox/flox^/*Prrx1*^Cre^ mice. We propose that combination of graded distribution of LRP1 and LRP1-mediated endocytic clearance and recycling of extracellular signaling molecules provides a novel mechanism for appropriate morphogen gradient formation to ensure that bones and joints form correctly. Investigations into the mechanisms underpinning formation of severe and persistent defects in skeletal elements caused by LRP1 loss are an essential for understanding the fundamental processes of morphogenesis, as well as the emergence of skeletal pathologies including DDH, osteoporosis and OA.

## Materials and Methods

### Mice

The *Lrp1^f^*^lox/flox^*/Prrx1*^Cre^ mice were generated by crossing the *Lrp1*^flox/flox^ (Strain 012604, the Jackson lab) and *Prrx1*^Cre^ (Strain 005584, the Jackson lab) mice. For post-natal analysis, 22 homozygotes, 14 heterozygotes and 21 wild-type littermates were examined for this study. For embryos, we examined 26 homozygotes, 12 heterozygotes and 24 wild-type littermates were examined for this study.

### Mouse genotyping

For genotyping, mice DNA was extracted from ear notches or tails using the RED ExtractN-Amp Tissue PCR Kit (SIGMA). For *Lrp1*^flox^, following primers were used; forward 5’- CATACCCTCTTCAAACCCCTTCCTG - 3’, reverse 5’-GCAAGCTCTCCTGCTCAGACCTGGA - 3’ with the following PCR conditions: 1 cycle of 94°C for 3 min, 35 cycles of 94°C for 30 s; 65°C for 30 s; 72°C for 30 s, ended with a final cycle of 72°C for 2 mins. For *Prrx1*^Cre^, following primers were used, forward 5’- GCTGCCACGACCAAGTGACAGCAA - 3’, reverse 5’- CAGTAGCCTCATCATCACTAGATG - 3’ with the following PCR conditions: 1 cycle of 94°C for 1 min, 40 cycles of 94°C for 30 s; 60°C for 30 s; 72°C for 1:30 mins; followed by a final cycle of 72°C for 5 mins.

### Immunohistochemistry

Slide sections were dewaxed and rehydrated by xylene and decreasing ethanol concentrations. Epitope unmasking was performed using a basic antigen retrieval reagent (R&D systems). The slides were immersed in the basic reagents and kept in a water bath for 10 minutes at 95°C. To block the endogenous peroxidase activity, 0.3% hydrogen peroxide was added to the slides and the slides were kept at 37°C for 15 minutes. Avidin/Biotin blocking kit (Vector Lab) was used to block endogenous biotin, biotin receptors and avidin. To block unspecific antibody binding, 10% goat serum was added to the slides and incubated for 3 hours at room temperature (RT). The primary antibody was then added to the slides and kept at 4°C overnight. The impress HRP Goat anti-Rabbit IgG polymer Detection Kit (Vector Lab) was used as a secondary antibody. One drop was used for each section, and the slides were kept at RT for 30 minutes. For signal enhancement, the VECTASTAIN ABC-HRP Kit (Vector Lab) was added to the slides and kept in the dark for 30 minutes. For signal visualisation, the DAB substrate kit (Vector Lab) was used to develop the brown signal. Fast green was used for counterstaining. Slides were dehydrated, cleared and mounted using increasing concentrations of ethanol, xylene and DPX (SIGMA), respectively. The primary antibodies used were as follows; rabbit monoclonal anti-LRP1 (1:200)(ab92544, Abcam), rabbit polyclonal anti-Sox9 (1:500)(AB5535, SIGMA), rabbit polyclonal anti-Wnt5a (1:200)(bs1948R, Bioss). The Rabbit IgG control antibody (I1000, Vector Lab) was used as isotype control. At least three mice/group were analysed for each staining.

### Microscopic Examination

For microscopic examination and imaging, the Nikon Eclipse Ci microscope with the DS-Fi2 high-definition colour camera head was used. All images were visualised using NIS-Elements imaging. The Zeiss Axio Scan.Z1 slide scanner system was used for the automated imaging. The resulting images were inspected using ZEN 3.0 (Blue edition), in which the image scale bar was added.

### Histology and staining

All samples were fixed with neutral buffered formalin overnight, then kept in 70% ethanol until processing. EDTA was used for bone decalcification with different incubation times based on the samples age. Sample processing was automated using the Leica ASP300 tissue processor (Leica Microsystems, UK). The Leica EG1150 H embedding station was used for sample embedding. Sample sectioning at 5 µm thickness was performed using the Leica RM2245 microtome. For H&E staining, slides were deparaffinised in xylene and rehydrated was performed using decreasing ethanol concentrations. For nucleus staining and counterstaining, slides were stained with haematoxylin and eosin, respectively, for 5 minutes. The slides were then dehydrated in increasing ethanol concentrations before clearing them in xylene followed by mounting. For Safranin O staining, sections were deparaffinised, rehydrated and stained with haematoxylin for 30 seconds followed by counterstaining with 2% fast green for 2 minutes. The sections were then dipped in acetic acid for 20 seconds before staining with safranin O for 8 minutes. At least three mice/group were analysed for each staining.

### Osteoblast isolation

Upper and lower limb bones from each mouse were chopped into small pieces, followed by collagenase digestion in a shaking water bath for 90 minutes at 37°C. After washing, the crushed bones were resuspended in 2ml of DMEM/F12 media with 10 % Fetal bovine serum (gibco, A384400-01) and then transferred to a T-25 culture flask. Clavaria osteoblasts were harvested the same way, but with collagenase digestion for 180 minutes. Osteoblasts were cultured and then subjected to SDS-PAGE followed by Western blot analysis.

### Micro-computed tomography (μCT)

For ex vivo high-resolution μCT imaging, all samples were fixed in buffered and then kept in 70% ethanol until processing. µCT scanning was performed using the Skyscan 1272 (SKYSCAN, Belgium) for all developmental stages. Based on the sample type and age, different imaging parameters were applied. For embryos, the parameters were as follows: 0.5 rotation step, 9μm isotropic resolution, 0.25 aluminium filter, 30 random movements and 4 average frames. For postnatal samples, the scanning parameters were as follows: 0.3 rotation step, 4.5μm isotropic resolution, 0.5 aluminium filter, 30 random movements and 2 average frames. The parameters for skull imaging were as follows: 0.5 rotation step, 9μm isotropic resolution, 0.5mm aluminium filter, 30 random movements and 2 average frames. Skyscan NRecon software was used to reconstruct the obtained images. Skyscan Data Viewer software was used to measure bone length and width. For trabecular bone, the Skyscan CT-analyser software was used to identify the trabecular bone in the tibia. CTvox software was used to visualise the obtained images in a 3D form.

For *in vivo* μCT, the University of Liverpool’s centre of preclinical imaging provided the imaging services using the Quantum GX-2 system (PerkinElmer, Inc. Waltham, MA). Whole-body scan images were acquired with the protocol FOV72, high speed, 8 sec x 3. High-resolution scans for upper and lower limbs were performed with the protocol FOV36, high resolution, 4 min. At least five mice/group were analysed for each staining.

### Locomotor activity monitoring

All mice were housed in a Digital Ventilated Cage (DVC®) rack, equipped with a home cage monitoring system capable of automatically measuring animal activity 24/7 (81). The DVC® rack is installed on a standard IVC rack (Tecniplast DGM500, Buguggiate, Italy) by adding sensing technologies externally to the cage, so that neither modifications nor intrusion occur in the home cage environment. Animal locomotion activity is monitored via a capacitance sensing technology by means of 12 contactless electrodes, uniformly distributed underneath the cage floor. The 6-week-old WT and cKO mice were monitored for 4 weeks and total distance and average speed of each mouse were measured.

### Whole-mount skeletal staining

The whole-mount skeletal staining was done following the Rigueur and Lyons’s protocol (82). Briefly, after dissection, adult mice skeletons were initially kept in 95% ethanol for 4 h before changing the solution and leaving in 95% ethanol overnight at RT. Then, skeletons mice were kept in acetone for two days at RT. Cartilage staining was performed by immersion in alcian blue for three days; then, mice underwent two changes of 95% ethanol for 4 hours and overnight for destaining. For pre-clearing, skeletons were kept in 1% KOH overnight at 4°C. Bone staining was performed by submersion in alizarin red for five days. Lastly, the final clearing was performed using 1% KOH prior to long-term storage in 100% glycerol.

### Double calcein labelling

Double calcein labelling was performed as described previously (83). Two intraperitoneal (IP) calcein injections (150 µl/mouse) were given to 5-6 weeks old mice 4 days before culling at an interval of two days. Mice were dissected, and the tibia bones were fixed in natural buffered formalin and kept in 70% ethanol until processing. The bones were dehydrated at 4°C with xylene and decreasing ethanol concentrations. Sample infiltration was performed under vacuum for seven days using a solution containing 88.99% Methyl Methacrylate (MMA), 10% dibutyl phthalate, 1% Perkadox 16 and 0.01% Novoscave. Teflon blocks filled with MMA were used for bone embedding. The blocks were kept at 30°C in a water bath to polymerise for 18 hours. After that, Historesin was used to attach the embedding rings. Sectioning at 5 µm was performed using the Leica RM2265 microtome. To analyse bone formation, sections were stained without de-plasticisation with 0.1% calcein blue for 3 minutes, dehydrated using different ethanol concentrations and cleared with xylene changes. Three mice/group were analysed.

### Tartrate-resistant acid phosphatase (TRAP) staining

Bone section de-plasticisation was performed by three changes of 2-methoxyethyl acetate (MEA). Section clearing and rehydration were accomplished by xylene and decreasing concentrations of ethanol. TRAP solution was prepared by dissolving naphthol ASTR-phosphate (1.4 mg/ml) and fast red (1.4 mg/ml) in a 0.2 M acetate buffer (pH 5.2) containing 100 mM sodium tartrate. Bone sections were kept in the TRAP solution for 2 hours at 37°C. Then bone sections were counterstained with 0.33 g/l aniline blue and 6 g/l phosphotungstic acid for 15 minutes. The slides were washed with distilled water and then cover slipped with Apathy’s serum. Three mice/group were analysed.

### Enzyme-linked immunosorbent assay (ELISA)

Purified human full-length LRP1 (10 nM in 100 µl of 50 mM Tris-HCl (pH 7.5)/150 mM NaCl/10 mM CaCl_2_, TNC) was coated overnight at 4°C onto microtiter plates (Corning, NY). Wells were blocked with 5% bovine serum albumin in TNC for 24 hours at 4 °C and washed in TNC containing 0.1% Brij-35 after this and each subsequent step. Wells were then incubated with various concentrations of Wnts in blocking solution for 30 min at RT. Bound proteins were detected using anti-Wnt3a antibody (ab219412, Abcam), anti-Wnt5a antibody (MAB6452, R&D systems) or anti-Wnt11 antibody (ab31962, Abcam) for 1 hour at RT and then with a secondary antibody coupled to horseradish peroxidase for 1 hour at RT. Hydrolysis of tetramethylbenzidine substrate (KPL, Gaithersburg, MA) was measured at 450 nm using a FLUOstar Omega (BMG Labtech). Mean values of technical duplicate were normalized by subtracting the amount of recombinant protein bound to control well that was not coated with LRP1. Extrapolated *K_D,app_* values were estimated based on one-phase decay nonlinear fit analysis using GraphPad Prism 9.

### Monitoring exogenously added Wnt5a levels in the cell culture

For 0-24 hours incubation assay, WT and LRP1 KO MEFs, and human chondrocytes (5 x 10^3^/well) were grown in 24 well plate (pre-coated with 0.1% gelatin for overnight) until cells reach confluent. Cells were then incubated with DMEM/F12 for overnight. The medium was replaced with 0.5 ml of fresh DMEM/F12 containing polymyxin B (50 µg/ml), CT1746 (100 µM) and the protease inhibitor cocktail (1/1000) with or without 20 nM Wnt5a in the presence or absence of 500 nM RAP. After 0-24 hours, 0.5 ml of medium were collected, and the protein was precipitated with trichloroacetic acid and dissolved in 40 μl of 1x SDS-sample buffer (50 mM Tris-HCl pH 6.8, 10 mM dithiothreitol, 2% SDS and 10% glycerol). Cells were washed with phosphate-buffered saline (PBS) once and lysed with 100 μl (for MEFs) or 50 μl (for chondrocytes) of 2x SDS-sample buffer. 12.5 µl of cell lysate and 5 µl (for MEFs) or 10 µl (for chondrocytes) of medium samples were analysed by SDS-PAGE under reducing conditions and underwent Western blotting using anti-Wnt5a antibody (MAB6452, R&D systems). Immune signals for exogenously added Wnt5a in the medium and cell lysate were quantified using ImageJ. Average actin signal was taken as 1 and Wnt5a signal in the cell lysate was normalised against each actin signal. The amount of Wnt5a after 24-h incubation was expressed as a % of the amount of Wnt5a after 3 hours (for MEFs) or 1 hour (for chondrocyte) incubation. For 0-60 minutes incubation assay, WT and LRP1 KO MEFs (5 x 10^3^/well) were grown in 96 well plate (pre-coated with 0.1% gelatin for overnight) until cells reach confluent. Cells were then incubated with DMEM/F12 for overnight. The medium was replaced with 50 µl of fresh DMEM/F12 containing polymyxin B (50 µg/ml), CT1746 (100 µM) and the protease inhibitor cocktail (1/1000) with or without 20 nM Wnt5a. After 0-60 minutes h, medium was removed, cells were washed with PBS once and lysed with 30 μl of 2x SDS-sample buffer. 10 µl of cell lysate samples were analysed by SDS-PAGE as described as above. Wnt5a immune signals in the cell lysate were quantified and normalised as described above and the amount of Wnt5a before incubation was taken as 100%.

### Immunocytochemical localisation of Wnt5a and LRP1

WT and LRP1 KO MEFs were generated as described previously (50) and kindly provided by Professor Dudley Strickland (University of Maryland School of Medicine). Human normal chondrocytes were prepared as described previously (9). Cells were grown with DMEM/F12 containing 10% FBS on 8-well Lab-Tek chamber slides (Nunc Lab-Tek Chamber Slide System, Thermo Scientific) precoated with 0.1% gelatin in PBS overnight. Once reaching confluency, the cells were rested in serum free DMEM/F12 for 24 h. Cells were then incubated in DMEM/F12 containing polymyxin B (50 µg/ml), the broad-spectrum hydroxamate metalloproteinases inhibitor CT1746 (100 µM) and the protease inhibitor cocktail (a mixture of aprotinin, bestatin, E-64, leupeptin and pepstatin A, 1/1000)(SIGMA, P1860) with or without 20 nM Wnt5a in the presence or absence of 500 nM RAP for 3 h at 37 °C. Cells were washed with DMEM/F12 three times and fixed with 4% paraformaldehyde (PFA) in PBS for 10 min at RT. PFA was removed and cells were incubated with 100 mM Glycine for 3 min at RT. Cells were washed with PBS twice and incubated with 0.1% Sudan black in 70% ethanol for 2 minutes. Cells were washed with 70% ethanol twice and PBS once, and permeabilised with in TNC containing 0.1% Triton X-100 for 3 minutes at RT. Cells were incubated with 10% goat serum for 1 hour at RT followed by three times washing with PBS. Each sample was then incubated with anti-Wnt5a antibody (AF645, R&D systems) and anti-LRP1 antibody (ab92544, Abcam) for overnight at 4 °C. Cells were washed with PBS three times and further incubated with Alexa Fluor 488-conjugated anti-mouse IgG and Alexa Fluor 568-conjugated anti-rabbit IgG (Molecular Probes, Eugene, OR) for one hour at RT. Actin was stained with Actin-stain 670 phalloidin (Cell Signalling). Cells were washed with PBS five times and mounted with VECTASHEILD antifade mounting media containing DAPI (2BScientific, Oxford, UK). Immunofluorescence images were acquired using a Zeiss LSM 800 confocal microscope (Zeiss, Oberkochen, Germany). Images were processed using Zen 2.6 (blue edition, Zeiss). For the three-dimensional image visualisation, the delineated region of z-stack image was loaded into the Imaris 10.0.1 software (Bitplane, Belfast, Northern Ireland, UK). The software was used to generate new spots layer. In this layer, LRP1 (red), Wnt5a (green) and DAPI (blue) channels were separately segmented. Slice view was used to measure spot diameter using the line tool and quality above 4.57 and 11.2 were used for LRP1 and Wnt5a, respectively.

### Xenopus laevis embryonic development model

All experiments were carried out in accordance with relevant laws and institutional guidelines at the University of East Anglia, with full ethical review and approval, compliant to UK Home Office regulations. Embryos were generated as described in (84). *Xenopus laevis* embryos were primed by injecting females with 100 U of PMSG into lymph sack 3 to 5 days before requiring the embryos. After 72 h, primed females were injected with 500 U of hCG into lymph sack to induce ovulation. Eggs were collected manually and fertilized in vitro. Embryos were de-jelled in 2% cysteine (w/v) solution until they pack closely together incubated at 18°C and microinjected in 3% Ficoll into 1 cell at the 2-4 cell stage into the dorsal marginal zone. Embryos were further incubated at 13° to 23°C until the required developmental stage was reached with observation and regular changing of 0.1x MMR. Embryo staging is according to Nieuwkoop and Faber (NF) normal table of Xenopus development (85). Wholemount in situ hybridisation was carried out as previously described (86). Sense and antisense probes were synthesised for lrp1. GFP, Wnt5a, Wnt11 and mini-LRP1 capped RNA for injections was prepared using T7 and SP6 mMessage mMachine kit (ThermoFisher, AM1344). Translation blocking morpholino antisense oligonucleotides targeting lrp1 (5’-TGCCTTGATCCAGTTCTTGG-3’) was purchased from Gene Tools, LLC in USA. A standard control morpholino (5’-CCTCTTACCTCAGTTACAATTTATA-3’) obtained from GeneTools was used for control injections.

### Immunofluorescence tissue staining of LRP1, Wnt5a, Vangl2 and phospho-Vangl2

Dewaxing and antigen retrieval was performed as described above. The slides were incubated with 0.1% Sudan black in 70% ethanol for 2 minutes. The slides were then washed with 70% ethanol twice and once with PBS for Wnt5a, LRP1 and Vangl2, and Tris-buffered saline (TBS) for phospho-Vangl2. To permeabilise the tissue, the slides were incubated with 0.4% Triton X-100 in PBS for Wnt5a, LRP1 and Vangl2, and in TBS for phospho-Vangl2 for 10 minutes at RT. To block unspecific antibody binding, 10% goat serum was added to all slides and incubated for 1 hour at RT followed by additional 1 hour mouse blocking (MKB 2213, Vector labs) for Wnt5a and LRP1 detection. The slides were washed twice with PBS for Wnt5a, LRP1 and Vangl2, and with TBS for phospho-Vavgl2. Each slide was then incubated with anti-Wnt5a antibody (Sc-365370, Santacruz), anti-LRP1 antibody (ab92544, Abcam), anti-Vangl2 antibody (21492-1-AP Proteintech Europe) and anti-phospho-Vangl2-S79/S82/S84 antibody (SAB5701946, Merck Life Science UK Limited) for overnight at 4 °C. The slides were washed four times with PBS for Wnt5a, LRP1 and Vangl2, and with TBS for phospho-Vavgl2. The slides were further incubated with Alexa Fluor 488-conjugated anti-mouse IgG and Alexa Fluor 568-conjugated anti-rabbit IgG (Molecular Probes, Eugene, OR) for 1 hour at RT. The slides were washed five times with PBS for Wnt5a, LRP1 and Vangl2, and with TBS for phospho-Vavgl2. The Slides were mounted and immunofluorescence images were acquired and processed as described above. Three mice/group were analysed.

### Statistical analysis

The GraphPad software program was used for statistical analysis. Significance was calculated using the unpaired Student’s t-test to compare each group and displayed on the statistical charts as * p<0.05, ** p<0.001, *** p<0.001 and **** p<0.0001. A two-way ANOVA was used for the comparison of cell-associated Wnt5a levels between WT and LRP1 KO MEFs after 5-30 min incubation. The data were represented as mean ±SD.

## Acknowledgments

We would like to thank Dr Kathryn Scott for helpful discussions during the writing of this manuscript. The Authors are grateful to the staff at the Biomedical Services, the Histology Units, the Centre for Preclinical Imaging and the Centre for Cell Imaging facilities provided by Liverpool Shared Research Facilities, Faculty of Health and Life Sciences, University of Liverpool.

This work was supported by the ministry of education of the Kingdom of Saudi Arabia (to M.A.), Libyan Ministry of Higher Education and Scientific Research (to A. ME. G.), Qatar National Research Fund (to N. A. AM.), European Union’s Horizon 2020 research and innovation programme under the Marie Skłodowska-Curie grant agreement (860635 to M. A. and A. K.), BBSRC Grant (BB/T00715X/1 to S. K. M. and G. N. W.), Versus Arthritis Career Development Fellowship (21447 to K.Y.) and Versus Arthritis Bridging Fellowship (23137 to K.Y.).

## Author Contributions

M Alhashmi and AME Gremida performed the experiments, acquired the data and wrote the manuscript. SK Maharana, M Antonaci and A Kerr performed the *Xenopus* experiments. NA Maslamani and MM Meschis supported M Alhashmi and AME Gremida to acquire the data. K Liu, H Sutherland and P Wilson performed the in vivo experiments and acquired the data. P Clegg provided financial support for the project and edited the manuscript. GN Wheeler directed the *Xenopus* experiments. RJ van’t Hof directed the µCT analysis. G Bou-Gharios designed the experiments, wrote and edited the manuscript. K Yamamoto performed the experiments, acquired the data, designed the experiments, wrote and edited the manuscript. All authors reviewed and revised the manuscript.

## Competing Interest Statement

The authors declare that there are no conflicts of interests.

**All supplementary materials can be found in a single supplementary information file.**

## References

1. Moestrup SK, Gliemann J, and Pallesen G. Distribution of the alpha 2-macroglobulin receptor/low density lipoprotein receptor-related protein in human tissues. Cell Tissue Res. 1992;269(3):375–82.

2. Zheng G, Bachinsky DR, Stamenkovic I, Strickland DK, Brown D, Andres G, et al. Organ distribution in rats of two members of the low-density lipoprotein receptor gene family, gp330 and LRP/alpha 2MR, and the receptor-associated protein (RAP). J Histochem Cytochem. 1994;42(4):531–42.

3. Strickland DK, Gonias SL, and Argraves WS. Diverse roles for the LDL receptor family. Trends Endocrinol Metab. 2002;13(2):66–74.

4. Bres EE, and Faissner A. Low Density Receptor-Related Protein 1 Interactions With the Extracellular Matrix: More Than Meets the Eye. Front Cell Dev Biol. 2019;7:31.

5. Troeberg L, Fushimi K, Khokha R, Emonard H, Ghosh P, and Nagase H. Calcium pentosan polysulfate is a multifaceted exosite inhibitor of aggrecanases. Faseb J. 2008;22(10):3515–24.

6. Scilabra SD, Troeberg L, Yamamoto K, Emonard H, Thogersen I, Enghild JJ, et al. Differential regulation of extracellular tissue inhibitor of metalloproteinases-3 levels by cell membrane-bound and shed low density lipoprotein receptor-related protein 1. J Biol Chem. 2013;288(1):332–42.

7. Yamamoto K, Okano H, Miyagawa W, Visse R, Shitomi Y, Santamaria S, et al. MMP-13 is constitutively produced in human chondrocytes and co-endocytosed with ADAMTS-5 and TIMP-3 by the endocytic receptor LRP1. Matrix Biol. 2016;56:57–73.

8. Yamamoto K, Owen K, Parker AE, Scilabra SD, Dudhia J, Strickland DK, et al. Low density lipoprotein receptor-related protein 1 (LRP1)-mediated endocytic clearance of a disintegrin and metalloproteinase with thrombospondin motifs-4 (ADAMTS-4): functional differences of non-catalytic domains of ADAMTS-4 and ADAMTS-5 in LRP1 binding. J Biol Chem. 2014;289(10):6462–74.

9. Yamamoto K, Scavenius C, Meschis MM, Gremida AME, Mogensen EH, Thogersen IB, et al. A top-down approach to uncover the hidden ligandome of low-density lipoprotein receptor-related protein 1 in cartilage. Matrix Biol. 2022;112:190–218.

10. Yamamoto K, Troeberg L, Scilabra SD, Pelosi M, Murphy CL, Strickland DK, et al. LRP-1-mediated endocytosis regulates extracellular activity of ADAMTS-5 in articular cartilage. Faseb J. 2013;27(2):511–21.

11. Yamamoto K, Santamaria S, Botkjaer KA, Dudhia J, Troeberg L, Itoh Y, et al. Inhibition of Shedding of Low-Density Lipoprotein Receptor-Related Protein 1 Reverses Cartilage Matrix Degradation in Osteoarthritis. Arthritis Rheumatol. 2017;69(6):1246–56.

12. Barnes H, Larsen B, Tyers M, and van Der Geer P. Tyrosine-phosphorylated low density lipoprotein receptor-related protein 1 (Lrp1) associates with the adaptor protein SHC in SRC-transformed cells. J Biol Chem. 2001;276(22):19119–25.

13. Boucher P, Liu P, Gotthardt M, Hiesberger T, Anderson RG, and Herz J. Platelet-derived growth factor mediates tyrosine phosphorylation of the cytoplasmic domain of the low Density lipoprotein receptor-related protein in caveolae. J Biol Chem. 2002;277(18):15507–13.

14. Loukinova E, Ranganathan S, Kuznetsov S, Gorlatova N, Migliorini MM, Loukinov D, et al. Platelet-derived growth factor (PDGF)-induced tyrosine phosphorylation of the low density lipoprotein receptor-related protein (LRP). Evidence for integrated co-receptor function betwenn LRP and the PDGF. J Biol Chem. 2002;277(18):15499–506.

15. Herz J, Clouthier DE, and Hammer RE. LDL receptor-related protein internalizes and degrades uPA-PAI-1 complexes and is essential for embryo implantation. Cell. 1992;71(3):411–21.

16. Nakajima C, Haffner P, Goerke SM, Zurhove K, Adelmann G, Frotscher M, et al. The lipoprotein receptor LRP1 modulates sphingosine-1-phosphate signaling and is essential for vascular development. Development. 2014;141(23):4513–25.

17. Hamlin AN, Basford JE, Jaeschke A, and Hui DY. LRP1 Protein Deficiency Exacerbates Palmitate-induced Steatosis and Toxicity in Hepatocytes. J Biol Chem. 2016;291(32):16610–9.

18. Ding Y, Xian X, Holland WL, Tsai S, and Herz J. Low-Density Lipoprotein Receptor-Related Protein-1 Protects Against Hepatic Insulin Resistance and Hepatic Steatosis. EBioMedicine. 2016;7:135–45.

19. Hofmann SM, Zhou L, Perez-Tilve D, Greer T, Grant E, Wancata L, et al. Adipocyte LDL receptor-related protein-1 expression modulates postprandial lipid transport and glucose homeostasis in mice. J Clin Invest. 2007;117(11):3271–82.

20. Hu L, Boesten LS, May P, Herz J, Bovenschen N, Huisman MV, et al. Macrophage low-density lipoprotein receptor-related protein deficiency enhances atherosclerosis in ApoE/LDLR double knockout mice. Arterioscler Thromb Vasc Biol. 2006;26(12):2710–5.

21. Overton CD, Yancey PG, Major AS, Linton MF, and Fazio S. Deletion of macrophage LDL receptor-related protein increases atherogenesis in the mouse. Circ Res. 2007;100(5):670–7.

22. Lin JI, Feinstein TN, Jha A, McCleary JT, Xu J, Arrigo AB, et al. Mutation of LRP1 in cardiac neural crest cells causes congenital heart defects by perturbing outflow lengthening. Commun Biol. 2020;3(1):312.

23. Boucher P, Gotthardt M, Li WP, Anderson RG, and Herz J. LRP: role in vascular wall integrity and protection from atherosclerosis. Science. 2003;300(5617):329–32.

24. Muratoglu SC, Belgrave S, Hampton B, Migliorini M, Coksaygan T, Chen L, et al. LRP1 protects the vasculature by regulating levels of connective tissue growth factor and HtrA1. Arterioscler Thromb Vasc Biol. 2013;33(9):2137–46.

25. Mao H, Lockyer P, Townley-Tilson WH, Xie L, and Pi X. LRP1 Regulates Retinal Angiogenesis by Inhibiting PARP-1 Activity and Endothelial Cell Proliferation. Arterioscler Thromb Vasc Biol. 2016;36(2):350–60.

26. Zhang JM, Au DT, Sawada H, Franklin MK, Moorleghen JJ, Howatt DA, et al. LRP1 protects against excessive superior mesenteric artery remodeling by modulating angiotensin II-mediated signaling. JCI Insight. 2023;8(2).

27. Mark PR, Murray SA, Yang T, Eby A, Lai A, Lu D, et al. Autosomal recessive LRP1-related syndrome featuring cardiopulmonary dysfunction, bone dysmorphology, and corneal clouding. Cold Spring Harb Mol Case Stud. 2022;8(6).

28. Bartelt A, Behler-Janbeck F, Beil FT, Koehne T, Muller B, Schmidt T, et al. Lrp1 in osteoblasts controls osteoclast activity and protects against osteoporosis by limiting PDGF-RANKL signaling. Bone Res. 2018;6:4.

29. Lu D, Li J, Liu H, Foxa GE, Weaver K, Li J, et al. LRP1 Suppresses Bone Resorption in Mice by Inhibiting the RANKL-Stimulated NF-kappaB and p38 Pathways During Osteoclastogenesis. J Bone Miner Res. 2018;33(10):1773–84.

30. Vi L, Baht GS, Soderblom EJ, Whetstone H, Wei Q, Furman B, et al. Macrophage cells secrete factors including LRP1 that orchestrate the rejuvenation of bone repair in mice. Nat Commun. 2018;9(1):5191.

31. Sims AM, Shephard N, Carter K, Doan T, Dowling A, Duncan EL, et al. Genetic analyses in a sample of individuals with high or low BMD shows association with multiple Wnt pathway genes. J Bone Miner Res. 2008;23(4):499–506.

32. Yan W, Zheng L, Xu X, Hao Z, Zhang Y, Lu J, et al. Heterozygous LRP1 deficiency causes developmental dysplasia of the hip by impairing triradiate chondrocytes differentiation due to inhibition of autophagy. Proc Natl Acad Sci U S A. 2022;119(37):e2203557119.

33. Dennis EP, Edwards SM, Jackson RM, Hartley CL, Tsompani D, Capulli M, et al. CRELD2 Is a Novel LRP1 Chaperone That Regulates Noncanonical WNT Signaling in Skeletal Development. J Bone Miner Res. 2020;35(8):1452–69.

34. Ono N, Balani DH, and Kronenberg HM. Stem and progenitor cells in skeletal development. Curr Top Dev Biol. 2019;133:1–24.

35. Cserjesi P, Lilly B, Bryson L, Wang Y, Sassoon DA, and Olson EN. MHox: a mesodermally restricted homeodomain protein that binds an essential site in the muscle creatine kinase enhancer. Development. 1992;115(4):1087–101.

36. Martin JF, and Olson EN. Identification of a prx1 limb enhancer. Genesis. 2000;26(4):225–9.

37. Logan M, Martin JF, Nagy A, Lobe C, Olson EN, and Tabin CJ. Expression of Cre Recombinase in the developing mouse limb bud driven by a Prxl enhancer. Genesis. 2002;33(2):77–80.

38. Kawanami A, Matsushita T, Chan YY, and Murakami S. Mice expressing GFP and CreER in osteochondro progenitor cells in the periosteum. Biochem Biophys Res Commun. 2009;386(3):477–82.

39. Henry SP, Jang CW, Deng JM, Zhang Z, Behringer RR, and de Crombrugghe B. Generation of aggrecan-CreERT2 knockin mice for inducible Cre activity in adult cartilage. Genesis. 2009;47(12):805–14.

40. Bi W, Deng JM, Zhang Z, Behringer RR, and de Crombrugghe B. Sox9 is required for cartilage formation. Nat Genet. 1999;22(1):85–9.

41. Akiyama H, Chaboissier MC, Martin JF, Schedl A, and de Crombrugghe B. The transcription factor Sox9 has essential roles in successive steps of the chondrocyte differentiation pathway and is required for expression of Sox5 and Sox6. Genes Dev. 2002;16(21):2813–28.

42. Haseeb A, Kc R, Angelozzi M, de Charleroy C, Rux D, Tower RJ, et al. SOX9 keeps growth plates and articular cartilage healthy by inhibiting chondrocyte dedifferentiation/osteoblastic redifferentiation. Proc Natl Acad Sci U S A. 2021;118(8).

43. Galea GL, Zein MR, Allen S, and Francis-West P. Making and shaping endochondral and intramembranous bones. Dev Dyn. 2021;250(3):414–49.

44. Green JL, Kuntz SG, and Sternberg PW. Ror receptor tyrosine kinases: orphans no more. Trends Cell Biol. 2008;18(11):536–44.

45. Gao B, Song H, Bishop K, Elliot G, Garrett L, English MA, et al. Wnt signaling gradients establish planar cell polarity by inducing Vangl2 phosphorylation through Ror2. Dev Cell. 2011;20(2):163–76.

46. Andre P, Wang Q, Wang N, Gao B, Schilit A, Halford MM, et al. The Wnt coreceptor Ryk regulates Wnt/planar cell polarity by modulating the degradation of the core planar cell polarity component Vangl2. J Biol Chem. 2012;287(53):44518–25.

47. Yamaguchi TP, Bradley A, McMahon AP, and Jones S. A Wnt5a pathway underlies outgrowth of multiple structures in the vertebrate embryo. Development. 1999;126(6):1211–23.

48. van Amerongen R, Fuerer C, Mizutani M, and Nusse R. Wnt5a can both activate and repress Wnt/beta-catenin signaling during mouse embryonic development. Dev Biol. 2012;369(1):101–14.

49. Mogensen EH, Poulsen ET, Thogersen IB, Yamamoto K, Bruel A, and Enghild JJ. The low-density lipoprotein receptor-related protein 1 (LRP1) interactome in the human cornea. Exp Eye Res. 2022;219:109081.

50. Willnow TE, and Herz J. Genetic deficiency in low density lipoprotein receptor-related protein confers cellular resistance to Pseudomonas exotoxin A. Evidence that this protein is required for uptake and degradation of multiple ligands. J Cell Sci. 1994;107 (Pt 3):719–26.

51. Kawata K, Kubota S, Eguchi T, Aoyama E, Moritani NH, Kondo S, et al. Role of LRP1 in transport of CCN2 protein in chondrocytes. Journal of cell science. 2012;125(Pt 12):2965–72.

52. Laatsch A, Panteli M, Sornsakrin M, Hoffzimmer B, Grewal T, and Heeren J. Low density lipoprotein receptor-related protein 1 dependent endosomal trapping and recycling of apolipoprotein E. PLoS One. 2012;7(1):e29385.

53. Croy JE, Shin WD, Knauer MF, Knauer DJ, and Komives EA. All three LDL receptor homology regions of the LDL receptor-related protein bind multiple ligands. Biochemistry. 2003;42(44):13049–57.

54. Meijer AB, Rohlena J, van der Zwaan C, van Zonneveld AJ, Boertjes RC, Lenting PJ, et al. Functional duplication of ligand-binding domains within low-density lipoprotein receptor-related protein for interaction with receptor associated protein, alpha2-macroglobulin, factor IXa and factor VIII. Biochim Biophys Acta. 2007;1774(6):714–22.

55. Neels JG, van Den Berg BM, Lookene A, Olivecrona G, Pannekoek H, and van Zonneveld AJ. The second and fourth cluster of class A cysteine-rich repeats of the low density lipoprotein receptor-related protein share ligand-binding properties. J Biol Chem. 1999;274(44):31305–11.

56. Scilabra SD, Yamamoto K, Pigoni M, Sakamoto K, Muller SA, Papadopoulou A, et al. Dissecting the interaction between tissue inhibitor of metalloproteinases-3 (TIMP-3) and low density lipoprotein receptor-related protein-1 (LRP-1): Development of a “TRAP” to increase levels of TIMP-3 in the tissue. Matrix Biol. 2017;59:69–79.

57. Niehrs C. The role of Xenopus developmental biology in unraveling Wnt signalling and antero-posterior axis formation. Dev Biol. 2022;482:1–6.

58. Wallingford JB, Fraser SE, and Harland RM. Convergent extension: the molecular control of polarized cell movement during embryonic development. Dev Cell. 2002;2(6):695–706.

59. Abu-Elmagd M, Garcia-Morales C, and Wheeler GN. Frizzled7 mediates canonical Wnt signaling in neural crest induction. Dev Biol. 2006;298(1):285–98.

60. Fisher M, James-Zorn C, Ponferrada V, Bell AJ, Sundararaj N, Segerdell E, et al. Xenbase: key features and resources of the Xenopus model organism knowledgebase. Genetics. 2023;224(1).

61. Bowes JB, Snyder KA, Segerdell E, Jarabek CJ, Azam K, Zorn AM, et al. Xenbase: gene expression and improved integration. Nucleic Acids Res. 2010;38(Database issue):D607–12.

62. Mikhailenko I, Battey FD, Migliorini M, Ruiz JF, Argraves K, Moayeri M, et al. Recognition of alpha 2-macroglobulin by the low density lipoprotein receptor-related protein requires the cooperation of two ligand binding cluster regions. J Biol Chem. 2001;276(42):39484–91.

63. Hartmann C, and Tabin CJ. Dual roles of Wnt signaling during chondrogenesis in the chicken limb. Development. 2000;127(14):3141–59.

64. Church V, Nohno T, Linker C, Marcelle C, and Francis-West P. Wnt regulation of chondrocyte differentiation. J Cell Sci. 2002;115(Pt 24):4809–18.

65. Feng D, Wang J, Yang W, Li J, Lin X, Zha F, et al. Regulation of Wnt/PCP signaling through p97/VCP-KBTBD7-mediated Vangl ubiquitination and endoplasmic reticulum-associated degradation. Sci Adv. 2021;7(20).

66. Sakai K, Hiripi L, Glumoff V, Brandau O, Eerola R, Vuorio E, et al. Stage-and tissue-specific expression of a Col2a1-Cre fusion gene in transgenic mice. Matrix Biol. 2001;19(8):761–7.

67. Roebroek AJ, Reekmans S, Lauwers A, Feyaerts N, Smeijers L, and Hartmann D. Mutant Lrp1 knock-in mice generated by recombinase-mediated cassette exchange reveal differential importance of the NPXY motifs in the intracellular domain of LRP1 for normal fetal development. Mol Cell Biol. 2006;26(2):605–16.

68. Sato A, Yamamoto H, Sakane H, Koyama H, and Kikuchi A. Wnt5a regulates distinct signalling pathways by binding to Frizzled2. Embo J. 2010;29(1):41–54.

69. Ohkawara B, Glinka A, and Niehrs C. Rspo3 binds syndecan 4 and induces Wnt/PCP signaling via clathrin-mediated endocytosis to promote morphogenesis. Dev Cell. 2011;20(3):303–14.

70. Wang Y, Chang H, Rattner A, and Nathans J. Frizzled Receptors in Development and Disease. Curr Top Dev Biol. 2016;117:113–39.

71. Church VL, and Francis-West P. Wnt signalling during limb development. Int J Dev Biol. 2002;46(7):927–36.

72. Hartmann C. A Wnt canon orchestrating osteoblastogenesis. Trends Cell Biol. 2006;16(3):151–8.

73. Pinson KI, Brennan J, Monkley S, Avery BJ, and Skarnes WC. An LDL-receptor-related protein mediates Wnt signalling in mice. Nature. 2000;407(6803):535-8.

74. Tamai K, Semenov M, Kato Y, Spokony R, Liu C, Katsuyama Y, et al. LDL-receptor-related proteins in Wnt signal transduction. Nature. 2000;407(6803):530-5.

75. Ren Q, Chen J, and Liu Y. LRP5 and LRP6 in Wnt Signaling: Similarity and Divergence. Front Cell Dev Biol. 2021;9:670960.

76. Zilberberg A, Yaniv A, and Gazit A. The low density lipoprotein receptor-1, LRP1, interacts with the human frizzled-1 (HFz1) and down-regulates the canonical Wnt signaling pathway. J Biol Chem. 2004;279(17):17535–42.

77. Terrand J, Bruban V, Zhou L, Gong W, El Asmar Z, May P, et al. LRP1 controls intracellular cholesterol storage and fatty acid synthesis through modulation of Wnt signaling. J Biol Chem. 2009;284(1):381–8.

78. Au DT, Migliorini M, Strickland DK, and Muratoglu SC. Macrophage LRP1 Promotes Diet-Induced Hepatic Inflammation and Metabolic Dysfunction by Modulating Wnt Signaling. Mediators Inflamm. 2018;2018:7902841.

79. Guo X, Day TF, Jiang X, Garrett-Beal L, Topol L, and Yang Y. Wnt/beta-catenin signaling is sufficient and necessary for synovial joint formation. Genes Dev. 2004;18(19):2404–17.

80. Shea CA, Rolfe RA, and Murphy P. The importance of foetal movement for co-ordinated cartilage and bone development in utero : clinical consequences and potential for therapy. Bone Joint Res. 2015;4(7):105–16.

81. Iannello F. Non-intrusive high throughput automated data collection from the home cage. Heliyon. 2019;5(4):e01454.

82. Rigueur D, and Lyons KM. Whole-mount skeletal staining. Methods Mol Biol. 2014;1130:113–21.

83. van ’t Hof RJ, Rose L, Bassonga E, and Daroszewska A. Open source software for semi-automated histomorphometry of bone resorption and formation parameters. Bone. 2017;99:69–79.

84. Hatch VL, Marin-Barba M, Moxon S, Ford CT, Ward NJ, Tomlinson ML, et al. The positive transcriptional elongation factor (P-TEFb) is required for neural crest specification. Dev Biol. 2016;416(2):361–72.

85. Faber J, and Nieuwkoop PD. Normal Table of Xenopus Laevis (Daudin): A Systematical & Chronological Survey of the Development from the Fertilized Egg till the End of Metamorphosis (1st ed.). Garland Publishing, *New York*. 1994.

86. Harrison M, Abu-Elmagd M, Grocott T, Yates C, Gavrilovic J, and Wheeler GN. Matrix metalloproteinase genes in Xenopus development. Dev Dyn. 2004;231(1):214–20.

