## Supplemental information for "Skeletal progenitor LRP1 deficiency causes severe and persistent skeletal defects with WNT/planar cell polarity dysregulation"

**Corresponding author:** Kazuhiro Yamamoto

Liverpool, L7 8TX, UK.

^*^These authors equally contributed to the work.

**This PDF file includes:**

Supporting text

Figures S1 to S4

Movies S1 and S2

Supporting Information Text

**Supplementary method**

*Establishment of cartilage Lrp1 conditional KO mice (Lrp1*^flox/flox^*/Acan*^CreERT2^*)*

The Lrp1/Acan mice were developed by mating the Acantm1(Cre-ERT2)Crm/J (strain 019148, the Jackson laboratory) and Lrp1flox (Strain 012604, the Jackson laboratory) mice. These *Lrp1*/*Acan* mice were tamoxifen-inducible. Therefore, upon tamoxifen's timely administration, the *Lrp1* gene was deleted in cartilaginous tissues in the generated offspring (Henry et al., 2009). Total 18 homozygotes and 28 wild-type littermates were examined for this study. These mice were genotyped the same way as the *Lrp1*/*Prrx1* mice. In these mice, the deletion of Exon 2 in the Lrp1 gene was also investigated. The following primers were used: forward 5'- TCTTCTACCCTATGAACCATTCCC - 3', and reverse 5'-TCTTGGCTCCTCCAGTGTTC - 3' with the following PCR conditions: 1 cycle of 94°C for 4 min, 40 cycles of 94°C for 30 s; 61°C for 30 s; 72°C for 1.5 min, followed by a final cycle of 72°C for 5 mins.

*Tamoxifen oral gavage*

Fifty mg of tamoxifen-free base (Sigma, Cat. No T5648) was suspended in 0.5 ml of 100% ethanol and fully dissolved in 4.5 ml of corn oil (SIGMA) by sonication (final 10 mg/ml)l. Similarly, the progesterone (Sigma, Cat. No P3972) was prepared following the same steps. The pregnant female was given either 100 or 200 µl of tamoxifen (1 or 2 mg/mice) and 50 µl progesterone (0.5 mg/mice) orally, gavage, twice in a timely manner to generate the experimental groups. Weight measurements were taken before the gavage administrations and mice termination.

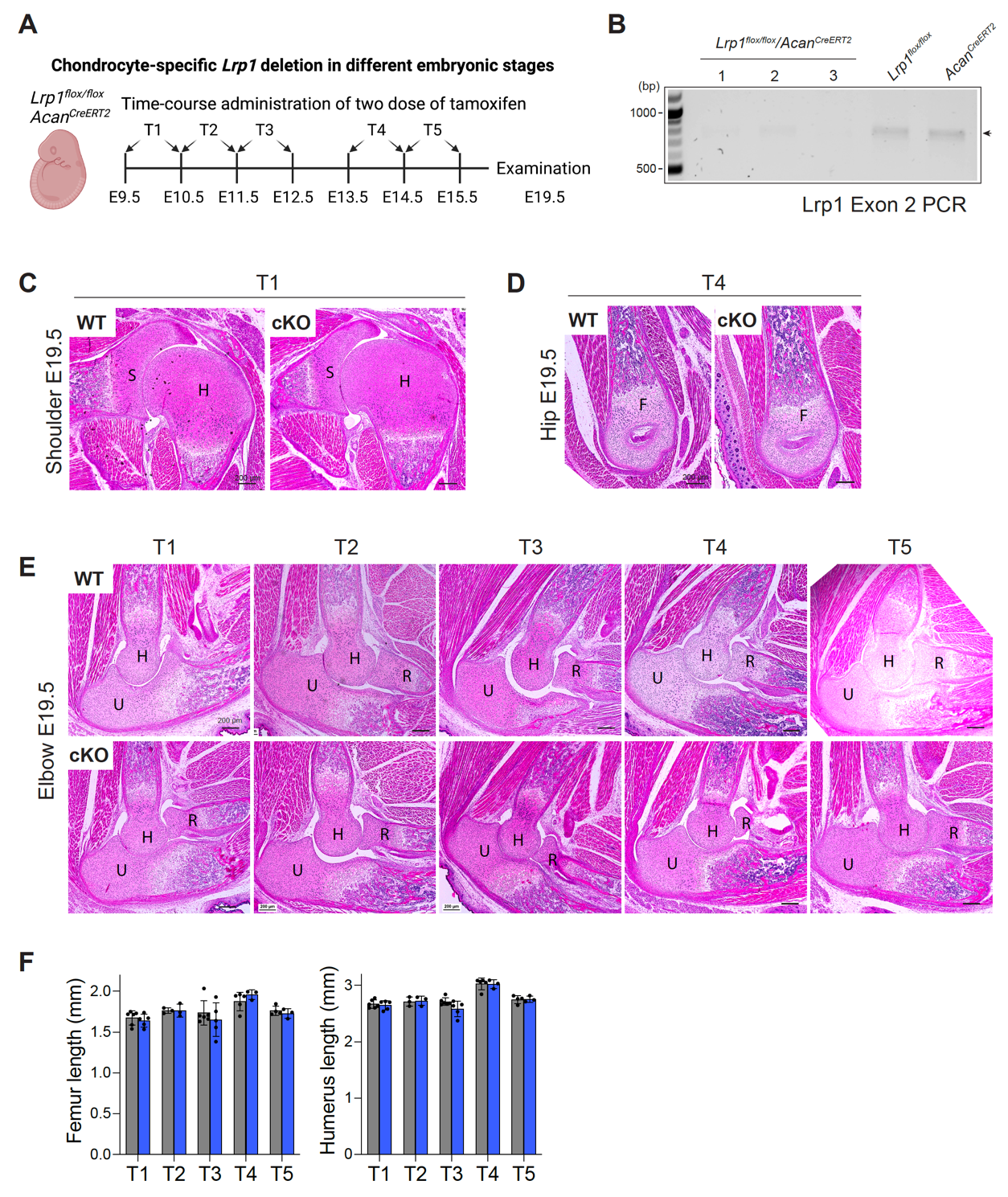

**Fig. S1. Conditional deletion of *Lrp1* in chondrocytes does not cause visible abnormality in early bone and joint formation.**

*A*, Schematic diagram showing the chondrocyte-specific *Lrp1* deletion (*Lrp1*^flox/flox^*/Acan*^CreERT2^) in mouse by two doses of tamoxifen gavage administration in the different embryonic stages. *B*, Representative agarose gel electrophoresis for *Lrp1*^flox/flox^*/Acan*^CreERT2^ genotyping to confirm the deletion of *Lrp1* *exon 2* gene. *C-F*, *Lrp1*^flox/flox^*/Acan*^CreERT2^ mice were given two doses of tamoxifen gavage in E9.5 and E10.5 (T1), E10.5 and E11.5 (T2), E11.5 and E12.5 (T3), E13.5 and E14.5 (T4), and E14.5 and E16.5 (T5). Representative images of H&E staining of E19.5 shoulder (*C*), hip (*D*) and elbow (*E*) of WT and cKO mice. Scale bar, 200 µm. S, scapula; H, humerus; U, ulna; R, radius; F, femur. *F*, Femur and humerus bone length of E19.5 WT and *Lrp1*^flox/flox^*/Acan*^CreERT2^ mice. Circles represent individual mice and bars show the mean ± *SD*.

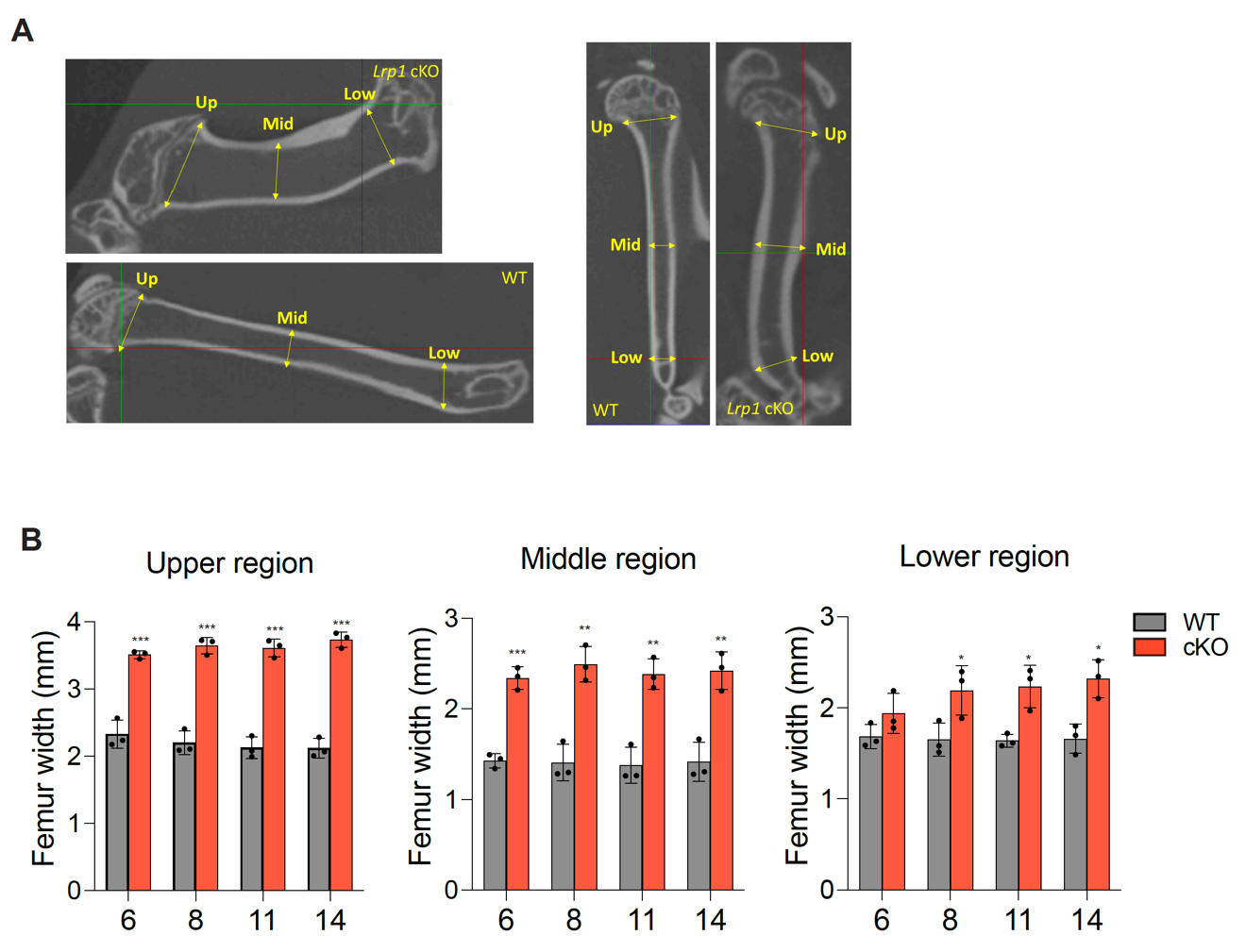

**Fig. S2. The upper, middle and lower region of femur width in WT and *Lrp1*^flox/flox^*/Prrx1*^Cre^ mice.**

*A*, Representative images for the bone width measurement on µCT images. The yellow arrows locate where the measurements were taken. (Up: Upper, Mid: Middle, Low: lower). *B*, Femur width at the upper, middle and lower regions of 6-14-week-old mice. Circles represent individual mice and bars show the mean ± *SD*. *, *p* < 0.05; **, *p* < 0.01; ***, *p* < 0.001; ****, *p* < 0.0001 by 2-tailed Student’s t test.

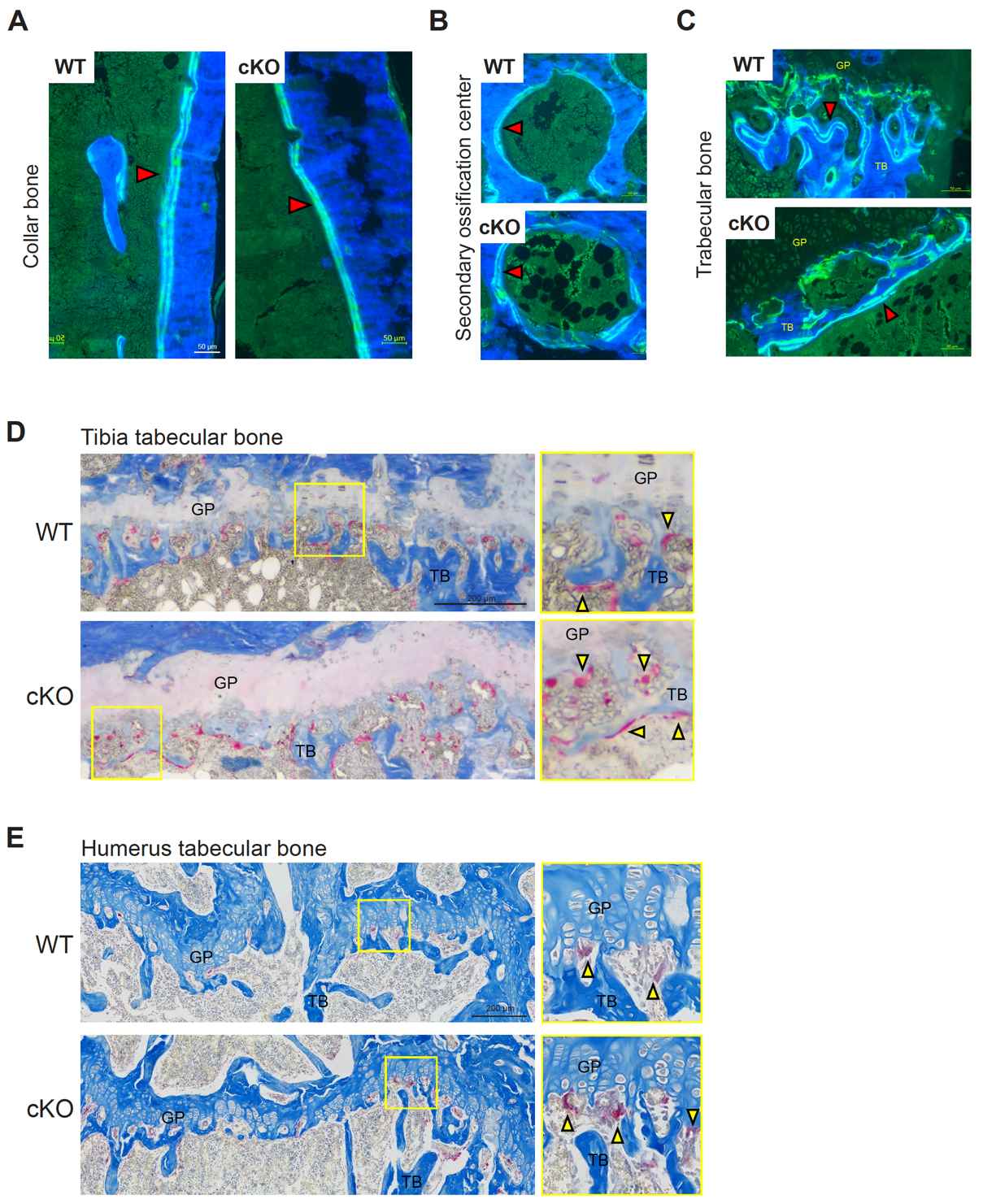

**Fig. S3. Comparable bone formation rate and an elevated number of osteoclasts in *Lrp1*^flox/flox^*/Prrx1*^Cre^ compared to WT mice.**

*A*-*C*, Representative images of fluorescent microscopy analysis of calcein-double stained histology sections of the collar bone (*A*), the secondary ossification center (*B*) and the trabecular bone (*C*) of 14-week-old WT and *Lrp1*^flox/flox^*/Prrx1*^Cre^ (cKO) mice. Scale bar, 50 µm. Red arrowheads indicate calcein double staining TB, trabecular bone; GP, growth plate. *D* and *E,* TRAP and aniline blue staining of 14-week-old tibia (*D*) and humerus (*E*) trabecular bones. Regions delineated by the squares in the left panels have been magnified in the right panels. Yellow arrowheads highlight osteoclast staining. Scale bar, 200 µm. TB, trabecular bone; GP, growth plate.

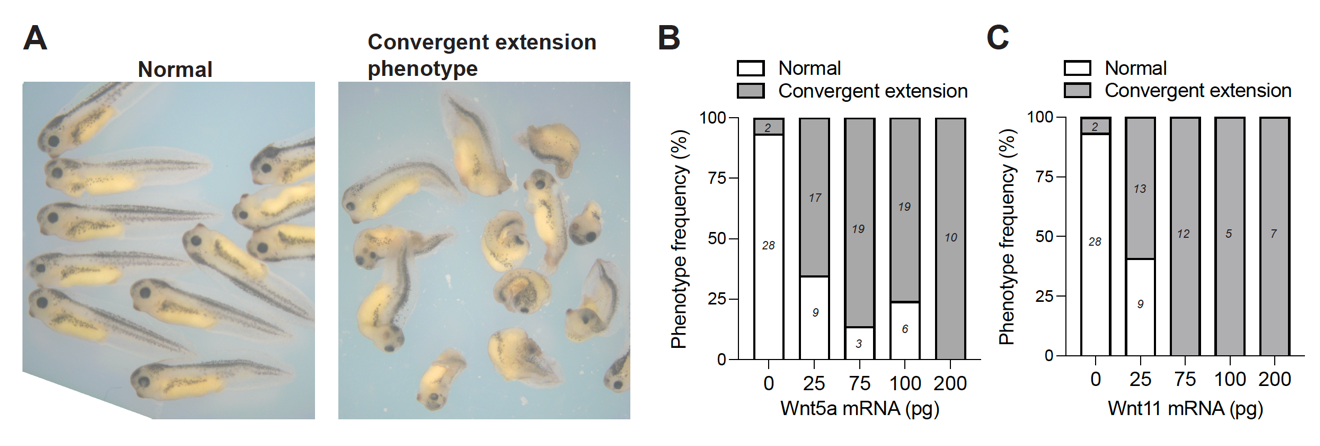

**Fig. S4. The WNT/PCP signalling controls convergent extension movements in the developing *Xenopus laevis* embryos.**

*A*, Representative images of normal Xenopus laevis tadpoles and those with disrupted convergent extension after overexpression of Wnt5a mRNA. *B and C*, Various amounts of mRNAs for *Wnt5a* (*B*) or *Wnt11* (*C*) were injected into 1 cell of the dorsal marginal zone of 4-cell stage embryos. Normal embryos and those with a convergent extension phenotype were counted after fixation at stage 35/36.

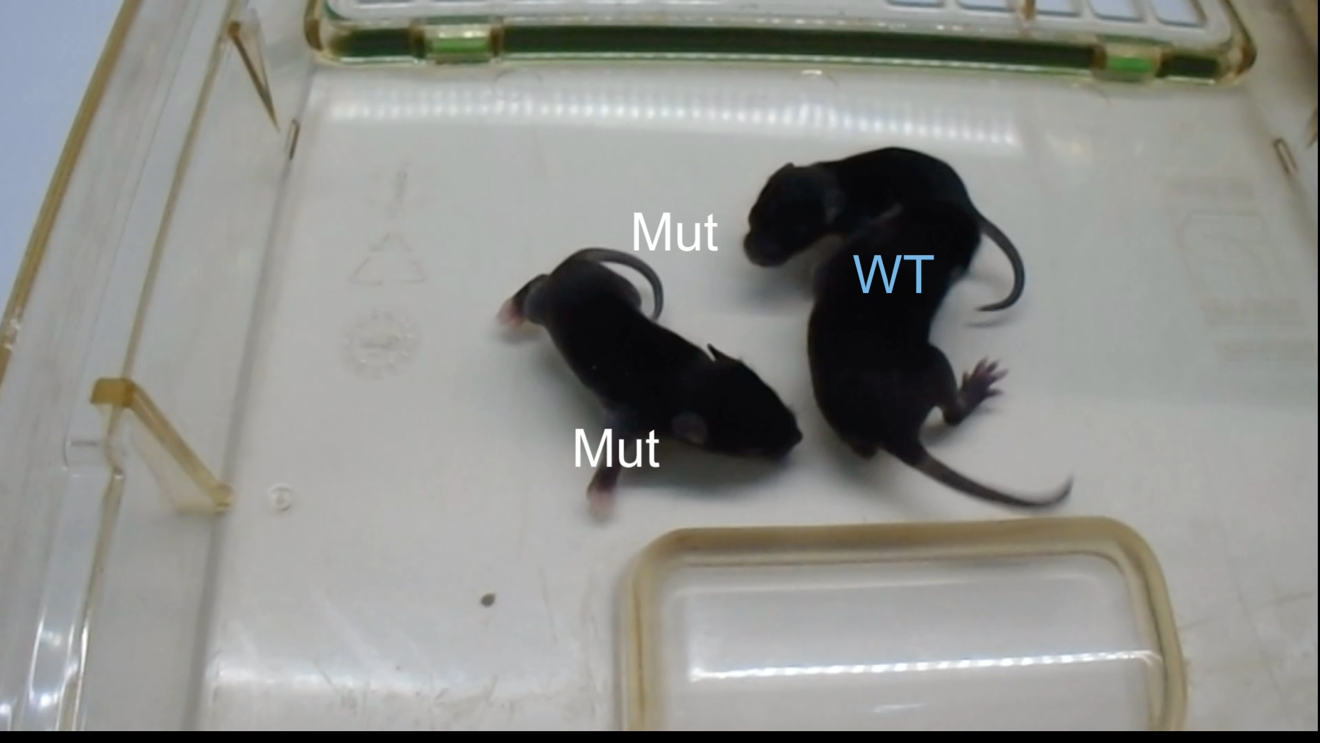

**Movie S1. Abnormal “crawling” gait of 2-week-old Lrp1 deficient mice.**

A short video showing abnormal gait of 2-week-old *Lrp1*^flox/flox^*/Prrx1*^Cre^ mice (Mut), which constantly show altered posture and impaired mobility. WT; wild-type littermate.

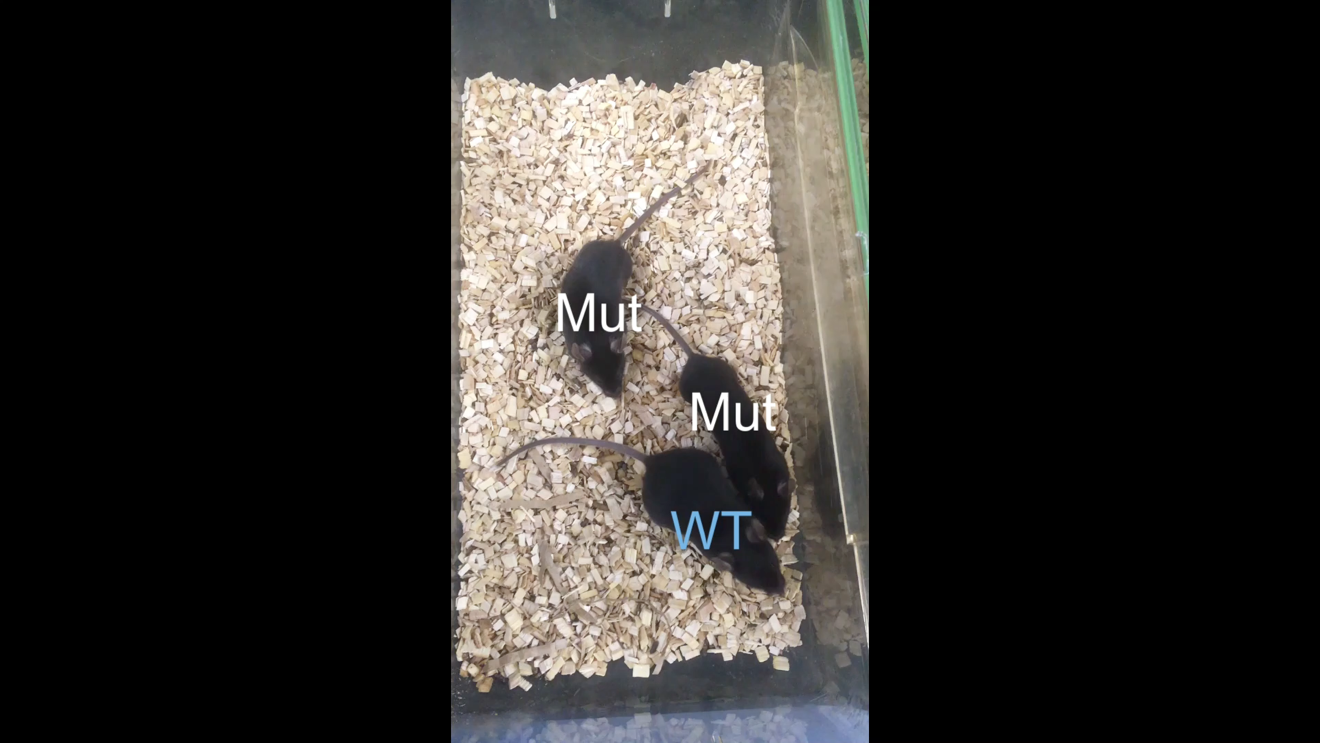

**Movie S2. Abnormal “crawling” gait of 13-week-old Lrp1 deficient mice.**

A short video showing abnormal gait of 14-week-old *Lrp1*^flox/flox^*/Prrx1*^Cre^ mice (Mut), which constantly show altered posture and impaired mobility. WT; wild-type littermate.
